# Structural analysis of the *C. elegans* dauer larval anterior sensilla by Focused Ion Beam-Scanning Electron Microscopy

**DOI:** 10.1101/2021.06.29.449894

**Authors:** Sebastian Britz, Sebastian Matthias Markert, Daniel Witvliet, Anna Maria Steyer, Sarah Tröger, Ben Mulcahy, Philip Kollmannsberger, Yannick Schwab, Mei Zhen, Christian Stigloher

## Abstract

At the end of the first larval stage, the nematode *Caenorhabditis elegans* developing in harsh environmental conditions is able to choose an alternative developmental path called the dauer diapause. Dauer larvae exhibit different physiology and behaviors from non-dauer larvae. Using focused ion beam scanning electron microscopy (FIB-SEM), we volumetrically reconstructed the anterior sensory apparatus of *C. elegans* dauer larvae with unprecedented precision. We provide a detailed description of some neurons, focusing on structural details that were unknown or unresolved by previously published studies. They include: 1) dauer-specific branches of the IL2 sensory neurons project into the periphery of anterior sensilla and motor or putative sensory neurons at the sub-lateral cords; 2) ciliated endings of URX sensory neurons are supported by both ILso and AMso socket cells near the amphid openings; 3) variability in amphid sensory dendrites among dauers; 4) somatic RIP interneurons maintain their projection into the pharyngeal nervous system. Our results support the notion that dauer larvae structurally expand their sensory system to facilitate searching for more favorable environments.

## 1 Introduction

The non-parasitic nematode *Caenorhabditis elegans* undergoes a rapid reproductive life cycle passing four larval stages (Brenner, 1974; Sulston and Horvitz, 1977). This underlies either self-fertilization by hermaphrodites or sexual reproduction involving males (Sulston and Horvitz, 1977). Harsh environmental conditions repress the reproductive life cycle and favor an alternative life cycle (Cassada and Russell, 1975), regulated by dauer pheromones (Golden and Riddle, 1982). Dauer larvae can survive for months without food (Klass and Hirsh, 1976). This is possible due to specific anatomical adaptations like a sealed mouth, a switch in metabolism to lipid storage, and protection from dehydration by specific changes in cuticles (Cassada and Russell, 1975; Popham and Webster, 1979; Albert and Riddle, 1983; Erkut et al., 2013). Consistently, dauer larvae are non-feeding, with inactive pharynx and reduced intestine (Cassada and Russell, 1975; Popham and Webster, 1979; Albert and Riddle, 1983).

The decision to enter dauer diapause is made in the first larva (L1) stage, triggered by poor environmental conditions including crowding, starvation and high temperature (Cassada and Russell, 1975; Golden and Riddle, 1984). When conditions do not provide enough food for the L1 larvae to become dauers, they arrest development and can survive without food for a few days (Johnson et al., 1984; Baugh and Sternberg, 2006). Arrested L1s do not have anatomical adaptions as pronounced as in dauer. For example, their mouth remains open therefore maintaining the ability to feed (Baugh and Sternberg, 2006; Baugh et al., 2009). Improvement of environmental conditions induces dauer exit and re-entry of reproductive development (Cassada and Russell, 1975; Golden and Riddle, 1982). As L1 is the only shared larval stage that is common between the reproductive and dauer developmental path before the dauer entry decision, the morphology of the L1 sensory apparatus is important as comparative complement.

Dauer larvae can perform, individually or as a group, a specific behavior called nictation. They raise and move their anterior body in a circular manner to increase the chance of getting attached to transport hosts and thereby being carried to a better environment, in a phoretic form of dispersal behavior (Sudhaus and Kiontke, 1996; Barrière and Félix, 2005; Lee et al., 2011; Félix and Duveau, 2012; Petersen et al., 2015; Yang et al., 2020). Anatomical, behavioral, and signaling adaptations exhibited by *C. elegans* dauers are reminiscent of parasitic nematodes at infectious stages (Reed and Wallace, 1965; Dieterich and Sommer, 2009; Ogawa et al., 2009). Nictation behavior was reported in insect-parasitic nematodes (Reed and Wallace, 1965; Gaugler and Campbell, 1993; Yang et al., 2020). Certain sensory neurons regulate dauer formation by an endocrine signaling system (Schackwitz et al., 1996) similar to that of parasitic nematodes (Ogawa et al., 2009).

The *C. elegans* nervous system consists of only a small number of neurons (Sulston and Horvitz, 1977) with similar morphologies and positions between individuals (White et al., 1986). It comprises two largely separated sub-systems – a somatic and pharyngeal nervous system – connected by two somatic interneurons (RIPL/R) (Ward et al., 1975; Albertson and Thomson, 1976; White et al., 1986). In non-dauers, RIPL/R’s anterior dendritic endings enter the pharyngeal basal lamina and form gap junctions with pharyngeal neurons (Ward et al., 1975; Albertson and Thomson, 1976). RIP neurons have been proposed to coordinate the activity between the pharyngeal and somatic nervous systems (Chalfie et al., 1985; Avery and Thomas, 1997). Whether RIPs maintain their projections to the dauer pharyngeal nervous system, which has become inactive (Cassada and Russell, 1975; Albert and Riddle, 1983), is unknown.

The somatic nervous system features multiple sensilla and non-sensilla associated sensory neurons that mediate a broad spectrum of sensory modalities from chemo-, mechano- to thermo-sensation that are important for development and dauer behaviors (Hedgecock and Russell, 1975; Albert and Riddle, 1983; Chalfie et al., 1985; Bargmann and Horvitz, 1991a, 1991b; Bretscher et al., 2011; Lee et al., 2011). Main sensory organs are located in the anterior tip, including six inner labial (IL), six outer labial (OL), four cephalic (CEP), and two amphid sensilla (Ward et al., 1975; White et al., 1986; Doroquez et al., 2014). IL, OL, CEP, and amphid neurons have ciliated sensory endings (Ward et al., 1975; Ware et al., 1975; Perkins et al., 1986; Doroquez et al., 2014). Dendritic endings of other sensory neurons (BAG, FLP, URX, and URY) project into this sensilla region, are ciliated, except for URY, and are not exposed to the external environment (Ward et al., 1975; White et al., 1986; Doroquez et al., 2014; Kazatskaya et al., 2020).

One of the most relevant neuron type for dauer behaviors are sensory IL2 neurons, which regulate nictation (Lee et al., 2011). They reside in the IL sensilla, each consisting of two sensory neurons (IL1 and IL2), enclosed by one socket (ILso) and one sheath (ILsh) cell (Ward et al., 1975; White et al., 1986; Doroquez et al., 2014). The socket cell anchors sensory neurons to hypodermis (Hyp), putatively for structural stability, and the sheath cell encloses neuron segments and may modulate perception (White et al., 1986; Doroquez et al., 2014). IL2’s dendrites show increased branching specifically in dauers revealing life-cycle depending phenotypic plasticity (Schroeder et al., 2013). In non-dauers, all six IL2 neurons have a single primary (1°) dendrite that projects anterior along the labial bundles (Schroeder et al., 2013; Androwski et al., 2020). In dauers, two dorsal and two ventral IL2 neurons (together called IL2Q) emanate in subsequent order of dendritic branches along 1° dendrites, some projecting between body wall muscle (BWM) and Hyp cell into the midline (*radial dendrites*) and then along the body wall (*body wall dendrites*), others (*midline dendrites)* projecting along the midline (Schroeder et al., 2013; Androwski et al., 2020), and additional branches (*dauer-specific 1° dendrites)* appear from IL2 soma (Schroeder et al., 2013). Two lateral IL2 neurons (called IL2L) exhibit a different dauer-specific dendritic pattern from IL2Q. The ends of their 1° dendrites emanate in subsequent order of short dendrites circumferentially reaching each other in some cases to form *crown-like* morphology (Schroeder et al., 2013). Ultrastructure and position in relation to anterior sensilla and further neurons in the sub-lateral cords of IL2 dendrites remain unclear.

Amphid and non-sensilla sensory neurons are critical for dauer development. Amphids consist of neurons that are important for dauer induction (ADF, ASG, ASI, ASJ, ASK) and recovery (ASJ) (Bargmann and Horvitz, 1991b; Schackwitz et al., 1996; Kim et al., 2009). Some sense dauer pheromones (ASI, ASK, ADL) (Macosko et al., 2009; Jang et al., 2012; Park et al., 2012) and temperature (AFD) (Liu et al., 2012). Other sensory neurons detect oxygen and carbon dioxide (BAG, URX) (Hallem and Sternberg, 2008; Bretscher et al., 2011; Hallem et al., 2011; Carrillo et al., 2013). Preferred foraging habitat of *C. elegans* is rotten fruits and plant stems, where carbon dioxide level and oxygen vary mostly due to the composting process (Gea et al., 2004; Blaxter and Denver, 2012; Félix and Duveau, 2012). Temperature regulates dauer induction and behaviors (Hedgecock and Russell, 1975; Golden and Riddle, 1984).

In non-dauers as well as dauers, anterior endings of lateral ILso (ILLso) cells have two branches that are enclosed by respective BAG and FLP neurons, while others (ILQso) do not feature such branches or interactions (Ward et al., 1975; Ware et al., 1975; Albert and Riddle, 1983; Perkins et al., 1986; Doroquez et al., 2014; Cebul et al., 2020). FLP neurons are highly branched along their dendrites and close to their cilia endings in adults (Albeg et al., 2011; Doroquez et al., 2014) while they only branch minimally in dauer larvae (Androwski et al., 2020). There was conflicting reporting on the sensory ending of URX neurons. URX were previously thought to be not ciliated (Doroquez et al., 2014), but recently reported to be ciliated in adults (Kazatskaya et al., 2020), and whether or not to be associated with ILLso cells (Doroquez et al., 2014; Cebul et al., 2020). Structures of URX dendritic endings in dauers have not being described.

In amphid sensilla, all sensory endings are enclosed by the amphid sheath cell (AMsh) (Ward et al., 1975; White et al., 1986; Doroquez et al., 2014). Amphid channel neurons (AM CNs: ASE, ADF, ASG, ASH, ASI, ASJ, ASK, ADL) expose ciliated endings through the amphid cuticle opening whereby the channel itself is formed by a socket cell (AMso). Other amphid sensilla sensory endings are enclosed by AMsh cells but do not enter the open channel (AWA, AWB, AWC, AFD) (Ward et al., 1975; White et al., 1986; Doroquez et al., 2014).

Regarding amphid sensilla, several structural differences were reported between dauers and non-dauers in previous studies (Ward et al., 1975; Ware et al., 1975; Albert and Riddle, 1983; White et al., 1986; Doroquez et al., 2014). The sensory endings of the temperature sensing neurons (AFD) were reported to be expanded its microvilli-like morphology in dauer larvae (Albert and Riddle, 1983) but it is important to note that the number of microvilli was only estimated in this study. In dauers, endings of AMsh cells were reported to be fused in approximately every second individual, and odor-sensing neurons (AWC) (Bargmann et al., 1993) expand their wing-like sensory endings more widely (Albert and Riddle, 1983; Procko et al., 2011).

Previous studies investigated the sensory structures of adult sensilla (Ward et al., 1975; Ware et al., 1975; White et al., 1986; Doroquez et al., 2014) and dauer sensilla with electron microscopy (EM) methods (Albert and Riddle, 1983). Classical chemical fixation is in particular limited in dauer due to their thick cuticles. High pressure freezing followed by careful freeze substitution (Weimer, 2006; Stigloher et al., 2011; Doroquez et al., 2014) now allows a near-to-native preservation of dauers (Schieber et al., 2017). Moreover, focused ion beam scanning electron microscopy (FIB-SEM) now enables the acquisition of comparably large volumes (Heymann et al., 2006) at very high Z-resolution (Briggman and Bock, 2012; Schieber et al., 2017). Combining the advantages of these emerging techniques, we applied them here to the anterior sensilla of dauer larvae. This allows us to obtain EM volumes with near-to-native ultrastructure at sufficient Z-resolution to reliably trace sensory neuron dendrites. Although our data sets provide more information, we focused on 3D reconstructions of anterior sensory neuron endings and their support cells and thereby describe unique structural features that have not been resolved in previous studies.

## 2 Materials and Methods

### 2.1 *C. elegans* maintenance and dauer induction

*Caenorhabditis elegans* N2 Bristol worms were maintained on 35 mm agar plates with nematode growth medium lite (Sun and Lambie, 1997) and *Escherichia coli* OP50 lawn at 20 °C (Brenner, 1974). Crowded populations where OP50 bacteria were still present were chunked onto 94 mm agar plates seeded with OP50 and cultured for seven days at 20 °C. Most worms developed into dauer stage due to overcrowding and starvation. Worms were washed off from plates with M9 buffer and treated with 1 % SDS solution for 10 min (Cassada and Russell, 1975). Five washing steps with M9 buffer were applied. To remove liquid, worms were centrifuged at 2,000g first. Worms were pipetted at the edge of a fresh unseeded agar plate for recovery and incubated for about 1-2 h. Dauer and some L1 larvae survived the SDS treatment and spread out over the plate. The agar areas with dead worms were removed with a spatula.

### 2.2 High pressure freezing, freeze substitution, and minimal resin embedding

SDS surviving dauer and L1 larvae were washed off with M9 buffer, which was then exchanged with 20 % bovine serum albumin in M9 buffer. Worms were pipetted into high pressure freezing planchettes with 100 μm recesses and high pressure frozen with an EM HPM100 machine (Leica) (Schieber et al., 2017). Worms were freeze substituted according to our published protocol in an EM AFS2 machine (Leica), infiltrated with Durcupan resin and then minimal resin embedded (Schieber et al., 2017).

### 2.3 FIB-SEM acquisition and image adjustment

Minimal resin embedded worms were further prepared for FIB-SEM imaging (Schieber et al., 2017). The data sets were acquired with a FIB-SEM Auriga 60 or Crossbeam 540 (Carl Zeiss Company) using ATLAS 3D software (part of Atlas5 from Fibics) by milling about 8 nm layers with the ion beam and imaging the block face with 5 nm pixel size. The data sets were acquired at 1.5 kV with the energy-selective back-scattered electron (EsB) detector (grid voltage 1100 V), at 700 pA. The anterior ends of three dauer larvae (*Dauer^E1^*, *Dauer^E2^*, *Dauer^E3^*) were imaged in transversal sections starting from posterior in the worm body orienting at specific anatomical landmarks of the worm. For *Dauer^E1^*, a larger body segment, starting at the amphid commissures was imaged. The anterior end of an L1 larva (*L1^E4^*) was imaged longitudinally.

Images were aligned using TrakEM2 in Fiji (Cardona et al., 2012; Schindelin et al., 2012). Image stack orientation regarding rotation (transversal), flipping, and stack recursion (longitudinal) as well as cropping, brightness, and contrast were adjusted either with TrakEM2 or Fiji. The image stack of *L1^E4^* was transformed first from a longitudinal to a transverse stack using the reslicing function of Fiji and then scaled to the respective pixel size by enlargement using a bilinear interpolation. In *Dauer^E2^* and *Dauer^E3^*, the VSNR denoising tool was used to reduce stationary artefacts (Fehrenbach et al., 2012). *L1^E4^* was denoised applying a Gaussian blur (sigma 1) in Fiji. The full volumes of *Dauer^E1^* and *Dauer^E3^* were prepared and exported with TrakEM2 and then uploaded to CATMAID (Saalfeld et al., 2009; Cardona et al., 2012).

For stage identification, we imaged a transverse section in the middle of the body of each worm to reveal cuticle, alae, and intestine morphology using the FIB-SEM. For additional investigation of the L1 larva, the remaining body fragments were re-embedded in Epon resin and longitudinally sectioned with an ultramicrotome. The sections were contrasted and imaged with a JSM-7500F SEM (JEOL) with our standard settings (Markert et al., 2017).

Images in figures and videos were adjusted and layouted as described in the **Supplementary Document**.

### 2.4 Larval stage and cell identification

We identified larval stage and sex of the three analyzed dauer larvae and the starved L1 larva by anatomical features listed in **Supplementary Table S1** and shown in **Supplementary Figure S1**. Left and right orientation were concluded from the known imaging direction and the other four body axes. This was validated for all data sets by analyzing the asymmetrical ventral nerve cord and asymmetry of certain neurons in the nerve ring in *Dauer^E1^*. In general, cells and other structures were identified with orientation provided by published maps (Ward et al., 1975; Albert and Riddle, 1983; White et al., 1986; Hall, 2008; Doroquez et al., 2014; Witvliet et al., 2020). All cells forming sensilla were identified in general by their position and morphology in the anterior end for all four data sets. Investigated neurons which are not part of sensilla (BAG, FLP, URX, URY, AVM, RIP) were identified by skeletal tracing up to the nerve ring region in *Dauer^E1^* first. Some of these neurons (BAG, URX, URY, RIP) with a characteristic morphology were identified in *Dauer^E2^*, *Dauer^E3^*, and *L1^E4^* by comparison of their anterior endings with those in *Dauer^E1^*. This was not reliably possible for others (FLP, AVM), as the morphology of their anterior endings was not characteristic in *Dauer^E1^*. RIP neurons endings were not part of the image volume of *Dauer^E2^*. As the dendrites of AM CNs (ASE, ADF, ASG, ASH, ASI, ASJ, ASK, ADL) could not be traced back to their nerve ring entry, they were identified for *Dauer^E1^* comparing their entry-order into AMsh cell as well as their location of the distal tips in the AMso cell with the literature (Ward et al., 1975; Albert and Riddle, 1983; Hall, 2008; Doroquez et al., 2014). These neurons could not be reliably identified in the other three data sets. Anterior projections of SAB and SAA in the ventral sub-lateral cords in *Dauer^E1^* could not be traced back to their nerve ring entry or somas and thus their identity not differentiated.

### 2.5 Volumetric reconstruction and cell tracing

Cells of interest in the most anterior 2,000 images of *Dauer^E1^* were volumetrically reconstructed (*Dauer^E1v^*) with the 3dmod package of the IMOD software (Kremer et al., 1996). The same cells were reconstructed in *Dauer^E2^* and *L1^E4^*. In general, cell contours were drawn manually at regular section intervals and contours of skipped sections interpolated with the 3dmod interpolator plugin. Some cells like sheath cells were encapsulating other cells or fusing with themselves, forming inward-facing surfaces which we did not reconstruct as the inner morphology was already defined by the ensheathed cells. Relevant cells in the data volume of *Dauer^E1^* were skeleton traced (*Dauer^E1s^*) using CATMAID. The tracing nodes were set into the centroid of the cells. This was performed for *Dauer^E3^* as well (*Dauer^E3s^*). In addition, cells of special interest were volumetrically reconstructed (*Dauer^E3v^*). Measurements were performed in this model as well. We traced some of the IL2Q dendrites retrogradely from the midlines to their 1° dendrite due to resolution limits.

Color code for cells was adapted from literature (Doroquez et al., 2014). Colors of left and right neurons were assigned different gradings for visual differentiation in some cases (AWA, AWB, AWC, FLP, IL2L, RIP). The cuticle of a larger volume of *Dauer^E1s^* shown in **Figure 1B** was automatically reconstructed with 3dmod applying *imodauto* function to an image stack rendering mask. This mask was created by application of filters and manual editing in Fiji, 3dmod, and GIMP (GIMP.org Team, 2021) functions. Automatically created contours were manually corrected in 3dmod. The cuticle of *Dauer^E1v^* shown in **Figure 2** was manually reconstructed as described above.

**Figure 1.**
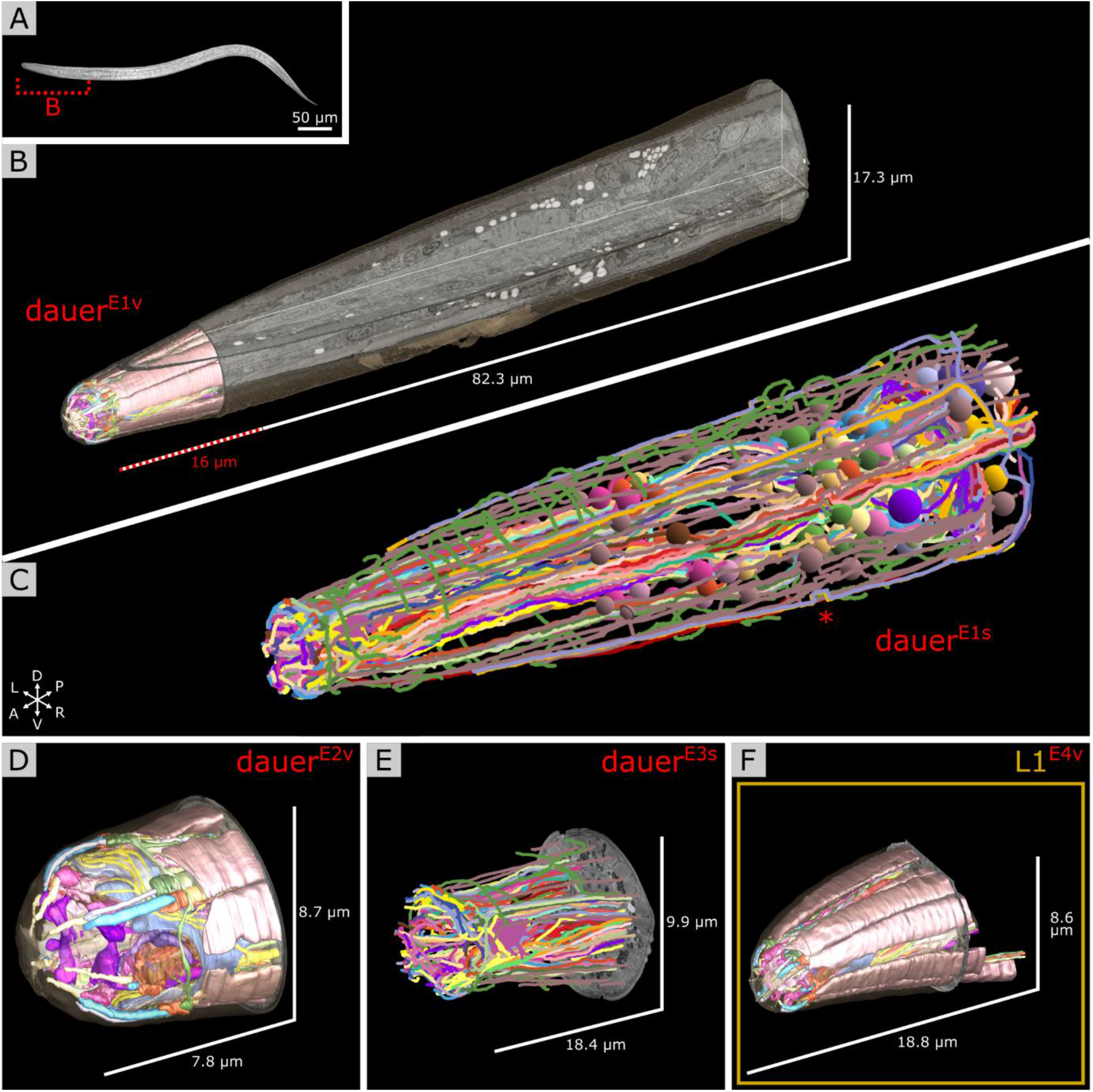
Electron microscopy reconstruction of the investigated head sensilla of *C. elegans* maintained under starving conditions. **(A)** Differential interference contrast image of a random dauer larva with a dashed line marking the estimated corresponding body segment from **B**. Anterior left. **(B)** Volumetric reconstruction of the head sensilla of one dauer larva in the context of the full FIB-SEM image volume used for cell tracing in **C**. X, Y, and Z image slices are shown. Reconstruction of the cuticle in the complete volume is shown. Scale refers to image volume. **(C)** Cell trajectories of the data set from **B** reconstructed by skeleton tracing. Color coded spheres label the identified somata. Images of a small segment are misaligned leading to a shift of tracing trajectories (asterisk). Data set is squeezed in Z by factor 0.625 due to incorrect settings. **(D)** Volumetric reconstruction of the head sensilla of a second dauer larva data set from a different individual. Skeleton tracing of the head sensilla of a third dauer larva. Data set is squeezed in Z by factor 0.625. Left is right in this data set. **(F)** Volumetric reconstruction of the head sensilla of a starving L1 larva.

**Figure 2.**
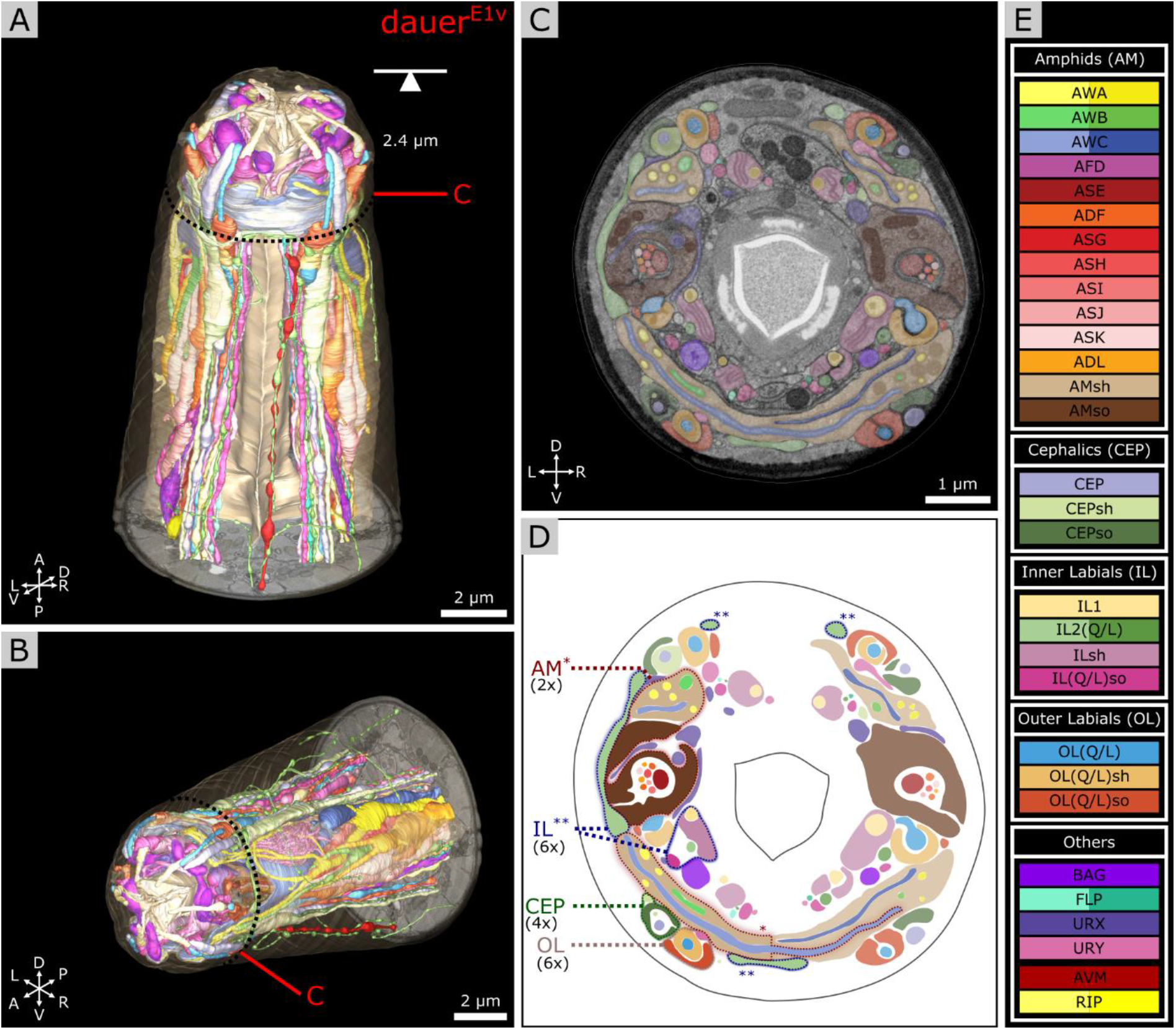
Volumetric reconstruction of 93 anterior cell endings including all sensory neurons and their support cells of a *elegans* dauer larva. **(A)** Ventral view of the volumetric reconstruction in context of body wall and buccal cuticle. **(B)** View from anterior-right side. **(C)** Transverse FIB-SEM section at 2.4 μm from anterior with cell labels according to color code (see **E**). **(D)** Scheme of cell labels from **C**. The four sensilla types called AM, IL, CEP, and OL are highlighted for one sensillum as an example each, indicating their location. Asterisks indicate associated parts respectively. **(E)** Cell names and color code (adapted from Doroquez et al., 2014). Cells are grouped by sensilla. AWA, AWB, AWC, lateral IL2, FLP, and RIP neurons are shown in different color gradients for left and right neurons for visual distinction. All cells shown are sensory neurons, except RIP neurons, and supporting socket (so) and sheath (sh) cells.

### 2.6 Measurements

Distances and lengths were measured with 3dmod by tracing the respective cell segment with an open contour and getting its length with *Edit > Contour > Info* for all four data sets (Kremer et al., 1996). The exact number of the dendritic endings of AWA was determined by drawing points on each end in 3dmod and counting the total number of points. Only endings distal to the ciliary base were counted. The length of the AWC wings was measured by a contour from end-to-end. In addition, the contour length was measured as flat contour applying *Edit > Object > Flatten*. The span of the AWC wings was measured end-to-end as follows. A point was set at each end and one in the center of the worm body in 3dmod. A snapshot facing the three points in parallel perspective (front view) in the 3dmod model view was taken. The snapshot was imported into Inkscape (Inkscape.org Team, 2021). The two endpoints were connected with the center-point each by two straight lines. The angle between the two lines was measured with the Inkscape measurement tool. This method was adapted from previous measurements of the wing span of AWC endings (Albert and Riddle, 1983). The number of AFD microvilli endings was determined in the same manner as the number of AWA endings.

## 3 Results

### 3.1 FIB-SEM data sets allow detailed 3D reconstruction of anterior sensilla

To get a near-to-native isotropic high-resolution 3D-insight into the anterior sensory apparatus of *C. elegans* dauer larvae we used high-pressure freezing followed by freeze substitution, minimal resin embedding, and FIB-SEM acquisition as previously described (Schieber et al. 2017).

We analyzed the anterior regions of sensilla of three dauer hermaphrodites by FIB-SEM image stacks, *Dauer^E1^*, *Dauer^E2^*, and *Dauer^E3^* (with *E* = EM data set). The number that follows the letter denotes different individuals, and the letter that follows the number denotes different reconstruction techniques (*v* = volumetric; *s* = skeletal). We applied volumetric reconstruction (*Dauer^E1v^*, *Dauer^E2v^*, *Dauer^E3v^*) (**Figure 1B,D**) or skeleton tracing (*Dauer^E1s^*, *Dauer^E3s^*) (**Figure 1C,E**). The *Dauer^E1v^* data set was restricted to the anterior sensilla region (**Figure 1B** and **Supplementary Video S1**) while *Dauer^E1s^* was traced to the amphid commissures (**Figure 1C** and **Supplementary Video S2**). Only relevant cells of *Dauer^E3^* were volumetrically reconstructed (*Dauer^E3v^*) (**Supplementary Figures S2B**, **S4D,E**, **S5C**). For comparative reasons we also acquired a data set of the anterior region of the sensilla of a starved L1 hermaphrodite that we reconstructed volumetrically (*L1^E4v^*) (**Figure 1F**). *Dauer^E1^*, *Dauer^E2^*, *Dauer^E3^* and *L1^E4^* larva were maintained under the same conditions. Features of these larvae and properties of these image stacks can be found in Methods and Supplementary information (**Supplementary Table S1** and **S2**; **Supplementary Figure S1)**.

### 3.2 Volumetric reconstruction reveals morphology of the anterior sensilla

For *Dauer^E1v^*, we present volumetric reconstructions of six IL, four CEP, six OL, two amphid sensilla, and sensory endings of URX, URY, FLP, and BAG (**Figure 2A,B**). 3D reconstructions of the endings of somatic sensory neuron AVM and interneurons (RIP) are included. The spatial organization of all sensilla and sensory neurons is shown in a transverse EM section (**Figure 2C,D**). Cells are labeled by the same color code used for the EM reconstruction of anterior sensilla of the adult *C. elegans* (Doroquez et al., 2014) (**Figure 2E**) to facilitate comparisons. The full reconstruction of *Dauer^E1v^* and additional details are shown in **Supplementary Video S1**.

Due to the improved Z-resolution and near-to-native sample preparation we could identify a number of features that have been previously either not reported, or misreported, which we now can present and clarify in detail.

### 3.3 Non-ciliated dendrites of IL2L project peripherally of all anterior sensilla

The *crown-like dendrites* of the inner labial sensilla IL2L cells are a specific feature of dauer larvae and their gross anatomy has been reported by fluorescence microscopy (Schroeder et al., 2013). Yet, the ultrastructural context of IL2L’s dauer *crown-like dendrites* and their location in relation to anterior sensilla is still unknown. We investigated and observed an overall similar anatomy for IL2L in all three dauer larvae. The location, morphology, and cellular structure of the *crown-like dendrites* of IL2L are shown for *Dauer^E1v^* (**Figure 3C**) and *Dauer^E3v^* (**Supplementary Figure S2A**).

**Figure 3.**
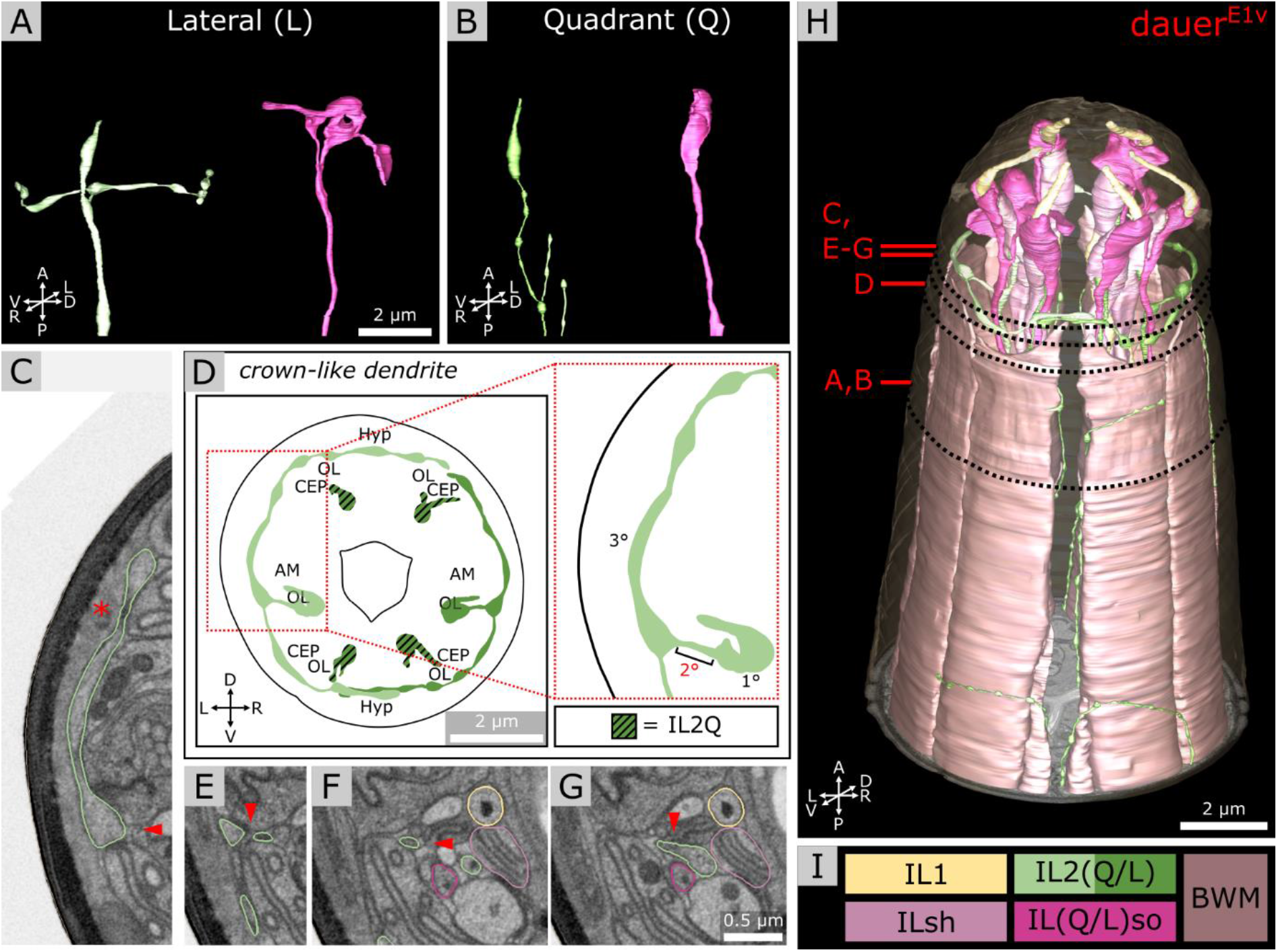
Morphology of dauer-specific *crown-like dendrites* of lateral IL2 endings. **(A)** Example of the morphology of IL2LL (panel left) and ILLso (panel right) cell endings. As an example, only left cells are shown. IL2LL has a *crown-like dendrite* projecting to dorsal and ventral side. ILLsoL shows two branches. **(B)** Example of the morphology of IL2Q (panel left) and ILQso (panel right) cell endings. Dorsal-left cells are shown as example. IL2QDL shows no *crown-like dendrite*, but several other dendritic branches are located more posterior. ILQsoDL shows no branches. **(D)** Schematic 2D projection of IL2 sensory endings including *crown-like dendrites* of a volume containing 5 μm from anterior of the model. IL2Q are graphicly hatched. **(C,E**-**G)** Four transverse FIB-SEM sections following the small 2° dendrite (arrowhead) from the 1° dendrite to the *crown-like dendrite* (which is a 3° dendrite). Asterisk in **C** indicates the Hyp syncytium, which together with the cuticle forms the only barrier to the environment for the *crown-like dendrite*. Panels are oriented on the same level in Y-dimension. **C** is on the same level as **Figure 2C**. **(H)** 3D reconstruction of all six IL sensilla in context with BWM cell endings. **(I)** Names and color code of cells shown.

For each IL2L neuron (**Figure 3A**, left), a small 2° dendrite was branching off the end of the 1° dendrite (**Figure 3C,E-G**), oriented in parallel to the anterior-lateral midline. These dendrites branched again to form a 3° dendrite, which we referred to as the *crown-like dendrites* (**Figure 3C,D,H**). The origin of the 2° dendrite was located posterior to the ciliary base. *Crown-like dendrites* had varicosities (**Figure 3A,D**) and in some cases (e.g. *Dauer^E3v^*) vesicular organelles (**Supplementary Figure S2A**). They projected to both, dorsal and ventral sides, each around the circumference of the worm head, mostly peripherally of amphid, CEP, and OL sensilla (**Figures 3C,D**, **5H**). *Crown-like dendrites* were peripherally only in contact with the Hyp syncytium which produces the cuticle (**Figure 3C**). We also observed *crown-like dendrites* to overlap in some cases (**Figure 3D,H** and **Supplementary Figure S2B**). There were minor differences in IL2L *crown-like dendrites* among dauer samples as they e.g. extended 4° dendrites in *Dauer^E3v^* (**Supplementary Figure S2B-D**) but not in other dauers (**Figure 3H**).

### 3.4 Dauer-specific branches of IL2Q reach SAB or SAA neurons

As described previously (Schroeder et al., 2013), we observed several dendritic branches on IL2Q dendrites (**Figure 4A** and **Supplementary Video S2**) following the body wall (*body wall dendrites*) (**Figure 4C,D**) or along the midline (*midline dendrites*) into both directions (**Figure 4B,G,I**). Both types originated mostly from a 2° *radial dendrite* which projected into the respective midline between BWM and the Hyp cell (**Figure 4C,F**). Some *body wall dendrites* originated from *midline dendrites* (**Figure 4B**). *Body wall dendrites* projected further from the midline along the body wall of the respective side, still between BWM and Hyp cell (**Figure 4D**). Some of them reached to SAB neurons in case of dorsal IL2Q dendrites (**Figure 4D,E**). Ventral IL2Q dendrites reached neurons in the sub-lateral cords as well, but here the identity of ventral SAB vs. SAA projections could not be clarified (**Figure 4G**). We were able to identify several IL2Q branches that were very clearly visible (**Figure 4F**). Some IL2Q *midline dendrites* ended in the midline (**Figure 4G**) and some were bundled with AVM. Dendritic IL2Q branches showed varicosities in most cases (**Figure 4D**).

**Figure 4.**
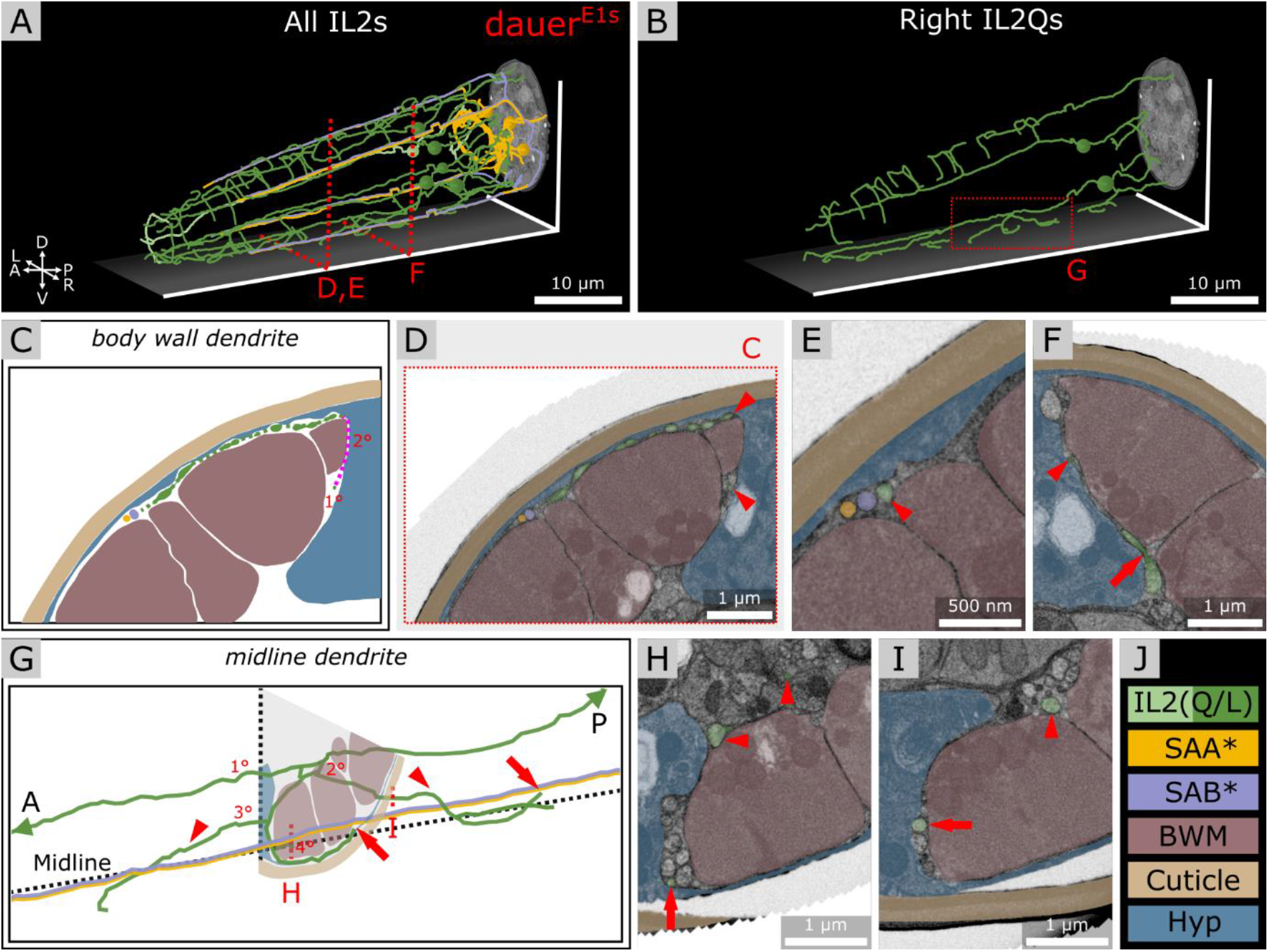
IL2Q neurons extend dauer-specific highly ordered branches all along their 1° dendrites, some of which come into close contact with SAA and SAB neurons. **(A)** Skeleton tracing of all six IL2 neurons, four SAA, and three SAB neurons in the anterior dauer larva. Spheres indicate the respective somata. Only IL2Q dendrites extend multiple branches along their length. IL2L neurons show anterior *crown-like dendrites* only. **(B)** Skeleton tracing of the right IL2Q neurons from **A**, as an example for IL2Q. **(C)** Scheme of an exemplary *body wall dendrite* of IL2QDL from **D** originating from a *radial dendrite* which is schematically delineated (magenta). **(D)** Transverse FIB-SEM section through the *body wall dendrite* of IL2QDL (arrowheads) used for scheme in **C**. **(E)** IL2QDL dendritic branch (arrowhead) from **D** reaching the SABD neuron (300 nm anterior to **D**). **(F)** Example of a *radial dendrite* of IL2QDR (arrowhead) one section before its branching point (arrow). **(G)** Scheme of IL2QVR dendrite branching indicated in **B** following the midline in both directions (arrowheads) and representing highly ordered branching up to 4° dendrite including *body wall dendrites* reaching neurons in the sub-lateral cord (arrows). Note that the identity of ventral SAB vs. SAA projections could not be clarified in our data set (*). **(H)** Transverse FIB-SEM section through IL2QVR (arrowheads) 4° dendrite (arrow). **(I)** Transverse FIB-SEM section through the IL2QVR (arrowhead) 2° dendrite (arrow) projecting posteriorly along the midline. **(K)** Names and color code of cells or structures shown.

### 3.5 Ciliated URX endings are supported by ILLso and AMso socket cells near amphid openings

The two branches of the ILLso cells interacted with non-IL sensilla cells (URX and BAG). One branch was facing into posterior direction originating from the socket ending (**Figure 3A**, right) and was enclosed by a certain part of the URX neuron (**Figure 5A**). This part of the URX neuron was then again enclosed by the AMso cell at the level where the AMso cell formed the channel of the AM CNs (**Figure 5B,D**). The AMso cell ended at the amphid opening of the cuticle and was enclosed there by OLLso cell (**Figure 5C**). URX endings were enclosed by AMso cells in all dauers. A similar but less extended ILLso-URX-AMso sandwich structure was observed across all dauers, in some cases (*Dauer^E2v^*) ILLso was in contact but not enclosed by URX. The described ILLso branch was missing in one case (*Dauer^E3s^*). URX endings were not enclosed by AMso cells in *L1^E4v^* but were enclosing respective branches of ILLso endings (**Supplementary Figure S3E,F**).

**Figure 5.**
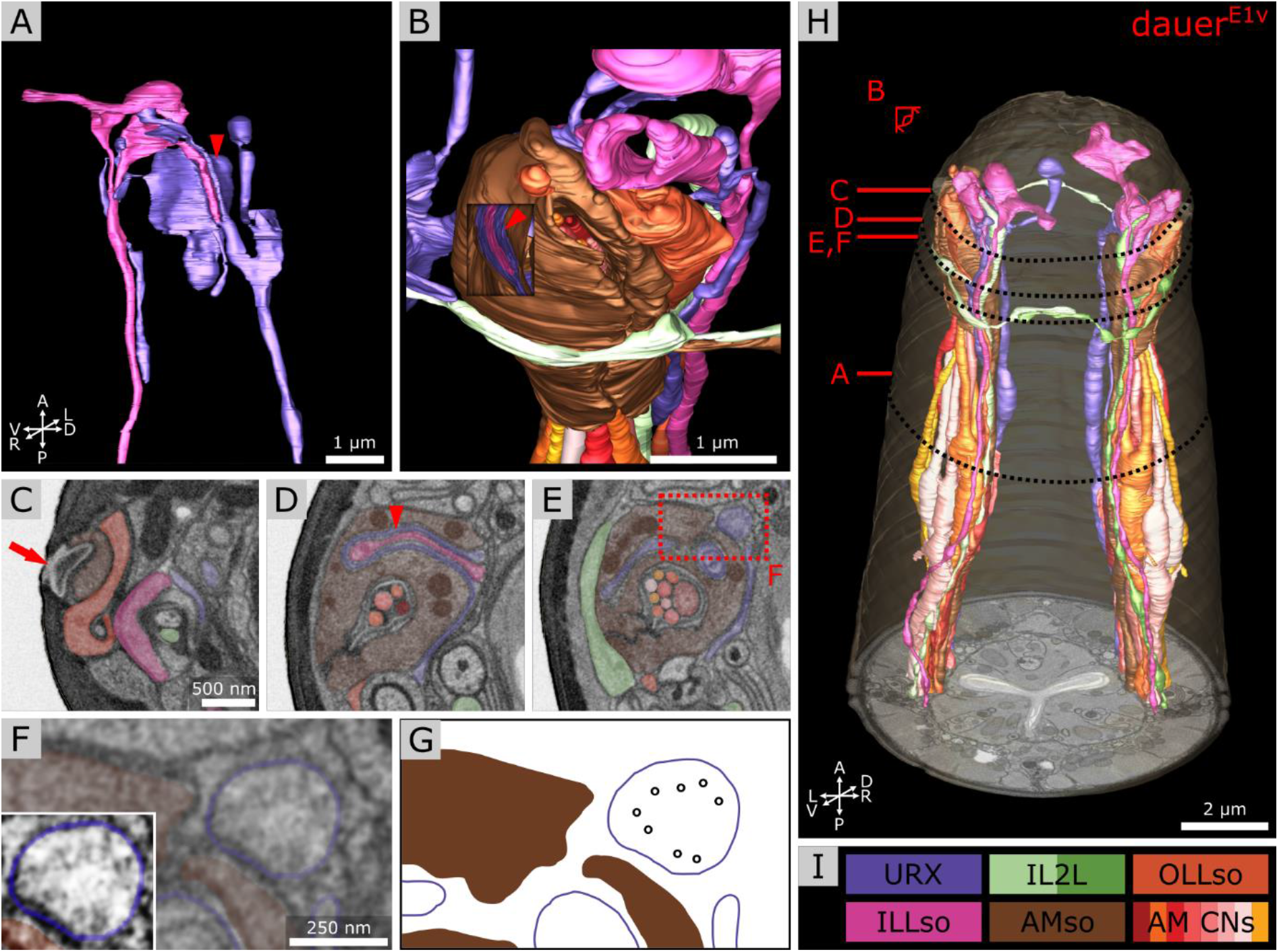
Anterior dendritic endings of dauer URX neurons are ciliated and extend distal structures to enclose ILLso and to be enclosed by AMso support cells. **(A)** Volumetric 3D reconstruction of URXL sensory ending enclosing a branch of ILLsoL (arrowhead). As an example, left side is shown. **(B)** Example of the spatial arrangement of URXL, ILLsoL, AMsoL, OLLsoL and IL2LL. Part of AMsoL is clipped to allow a view inside. AMsoL encloses the part of URXL which encloses the branch of ILLsoL shown in **A** (arrowhead). OLLsoL encloses AMsoL at the amphid opening. *Crown-like dendrite* of IL2LL is in direct contact with AMsoL. **(C)** Transverse FIB-SEM section at the amphid opening (arrow) in the cuticle, which marks the beginning of the amphid channel formed by AMsoL. AMsoL is enclosed by OLLsoL. **(D)** Transverse section at the position where AMsoL encloses URXL which encloses ILLsoL (arrowhead). **(E)** Transverse section at the position where AMsoL encloses URXL but without enclosing ILLsoL. The IL2LL branch passes AMsoL at the periphery. **(F)** Higher magnified part of **E** showing microtubules inside URXL. Inset shows same image with enhanced contrast. **(G)** Scheme indicating microtubules in **F**. **(H)** Overview of reconstruction of all mentioned cell endings. **(I)** Names and color code of cells shown.

Compared to the branch of ILLso described above, its second branch was originating from the cell rod and enclosed by BAG (**Supplementary Figure S3B,C**) which was facing into anterior direction (**Figure 3A**, right) just below the cuticle surface (**Supplementary Figure S3B,C**). We observed this in all three dauer larvae and to a less extent in *L1^E4v^* where BAG enclosed a swelling at the respective ILLso cell rod posterior to its socket.

We observed microtubules in the URX endings, suggesting that those are ciliated (**Figure 5F,G**). We also observed convoluted whorls of membranes and large vesicles, enclosed either by URX or the AMso cell (**Supplementary Figure S3A**). We found similar membranous structures enclosed by BAG or ILLso cells of which some originated from Hyp cells or URX (**Supplementary Figure S3B**). However, we could not determine their origin exactly.

Each FLP neuron had one 1° dendrite following the lateral labial bundle to anterior of which each extended two 2° dendrites following dorsal or ventral labial bundles respectively (**Supplementary Figure S3D**). These anterior projections were almost unbranched. Their endings neither appeared to be branched in *Dauer^E1v^* (**Supplementary Figure S3C**). We could not find any cilia microtubules in FLP. During the investigation of the ventral midline of *Dauer^E1s^*, we followed the AVM neuron and found its end anterior in the dauer at approximately 5 μm from the tip of the worm (**Supplementary Figure S3C,D**) reaching as far as the respective BWM cell.

### 3.6 Volumetric reconstruction of dauer amphids at single microvilli resolution

We determined the exact number of dendritic endings at the sensory tip of AWA neurons in dauer (**Table 1** and **Figure 6A**). Branches of AWA neurons in *Dauer^E3v^* did widely overlap (**Supplementary Figure S4D**) which was not the case in *Dauer^E1v^* and just partly in *Dauer^E2v^*. We also counted AWA endings in *L1^E4v^* (**Table 1**). One AWA neuron showed a wing-like expansion in one branch in *L1^E4v^* (**Supplementary Figure S4F**).

**Table 1.** Measurements of amphid neuron endings. Number of AWA dendritic endings. Span and end-to-end (contour) length (in 2D and 3D) of AWC neuron wings. 2D means length of flat contour. Number of AFD microvilli endings.

| Results per neuron | Neuron | Dauer<sup>E1</sup> | Dauer<sup>E2</sup> | Dauer<sup>E3</sup> | L1<sup>E4</sup> |
| --- | --- | --- | --- | --- | --- |
| Number of dendritic endings | AWAL | 14 | 15 | 24 | 8 |
|  | AWAR | 12 | 20 | 17 | 4 |
| Span of wings end-to-end [°] | AWCL | 294.57 | 298.72 | 342.95 | 52.24 |
|  | AWCR | 288.88 | 238.32 | 281.23 | 51.36 |
| Contour length of wings end-to-end [µm] | 2D AWCL | 12.7 | 14.6 | 15.6 | 2.5 |
|  | AWCR | 11.9 | 11.4 | 12.6 | 2.9 |
|  | 3D AWCL | 14.1 | 15.2 | 18.7 | 3.1 |
|  | AWCR | 14.4 | 12.5 | 13.2 | 3.1 |
| Number of microvilli endings | AFDL | 106 | 102<br>with 25 cut off | 85 | 37 |
|  | AFDR | 112 | 84<br>with 28 cut off | 88 | 37 |

**Figure 6.**
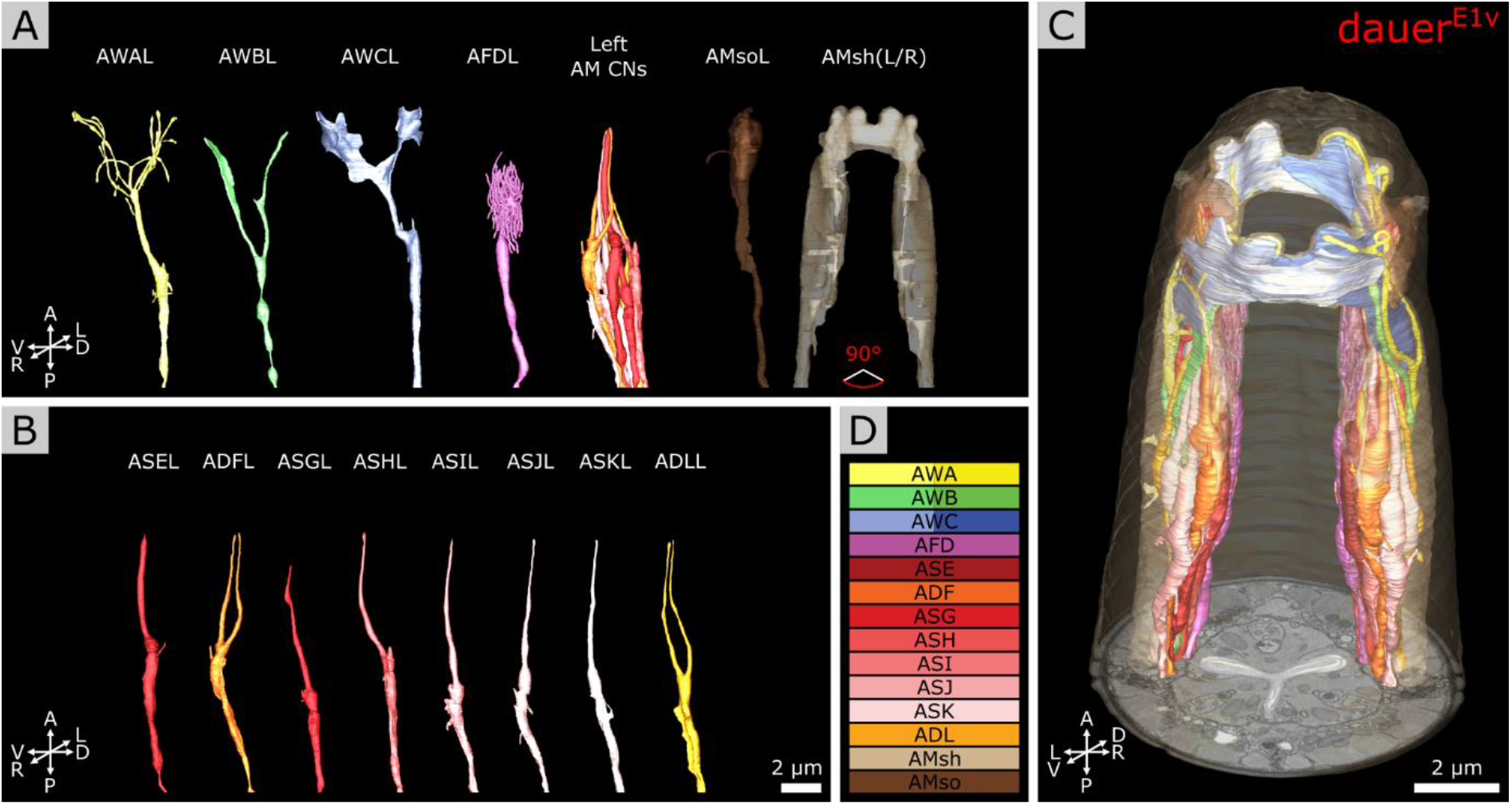
3D volumetric reconstruction of the anterior sensory endings of amphid neurons in dauer. **(A)** Models of sensory endings of all twelve left amphid neurons, of which eight are amphid channel neurons shown as a group (AM CNs), and models of the endings of their two support cells AMsoL and AMshL, where the latter is fused with AMshR. Left cells are shown as an example. **(B)** Models of sensory endings of all eight left amphid channel neurons from **A**. ADFL and ADLL are double ciliated while all others are single ciliated. **(C)** Models of sensory endings of all amphid neurons and their support cells. **(D)** Names and color code of cells shown.

Notably, we observed variability in the branching patterns of AWB as the two branches of the AWB neuron endings extended into dorsal and ventral direction in dauer (**Figure 6A**) while in one case of AWBL of individual *Dauer^E3v^* both branches extended into ventral direction only (**Supplementary Figure S4E**).

The prominent wing-like endings of AWC neurons were Y-shaped (**Figure 6A**) in dauers. The AWC wings overlapped on the dorsal and ventral sides (**Figure 6C**) and intertwined in some cases. We measured the span and length of AWC wings from end-to-end across all dauer larvae (**Table 1**). The wing of AWCR had a smaller span in *Dauer^E2v^* (**Table 1**). This wing fused with itself enclosing AMsh cytoplasm at two positions (**Supplementary Figure S4A,B,C**) which was only seen in this data set. The AWC wings in *L1^E4v^* were less expanded and did not overlap (**Table 1**).

We determined the exact number of AFD microvilli endings (**Figure 6A** and **Table 1**). In *Dauer^E2v^*, some posterior microvilli might be missing, because they were cut off in the data set. Therefore, we provide here the number of cut off microvilli in the reduced data set in addition, being aware that this is an underestimate of the total number (**Table 1**). We also counted AFD microvilli endings in *L1^E4v^* (**Table 1**).

### 3.7 RIP neuron maintains their projection into the pharynx in dauers

We found that in dauers, RIP neurons (**Figure 7A**) maintain their projections to the pharyngeal nervous system (**Figure 7F**). They entered through the pharyngeal basal lamina (**Figure 7B**), similarly in *L1^E4v^* (**Supplementary Figure S5E**). Posterior to the position where the endings migrated into the pharynx, they featured a bouton containing electron dense vesicles in both *Dauer^E1v^* and *Dauer^E3v^* (**Figure 7E** and **Supplementary Figure S5A,C**) and to a lesser extent in *L1^E4v^* (**Supplementary Figure S5D**). Inside the pharyngeal nervous system, RIP endings of *Dauer^E1v^* and *Dauer^E3v^* formed such bouton-like shape as well (**Figure 7F** and **Supplementary Figure S5C**) filled with electron dense vesicles (**Figure 7C**), which were also present in *L1^E4v^* but again to a lesser extent (**Supplementary Figure S5E**). The bouton of RIPL located posterior to the entry into the pharynx in *Dauer^E3v^* was enclosed by the respective BWM cell (**Supplementary Figure S5A,C**). We observed a readily identifiable active zone-like structure inside the bouton of RIPL of *Dauer^E3v^* opposing the BWM, which suggests that it is a neuromuscular junction (**Supplementary Figure S5B**).

**Figure 7.**
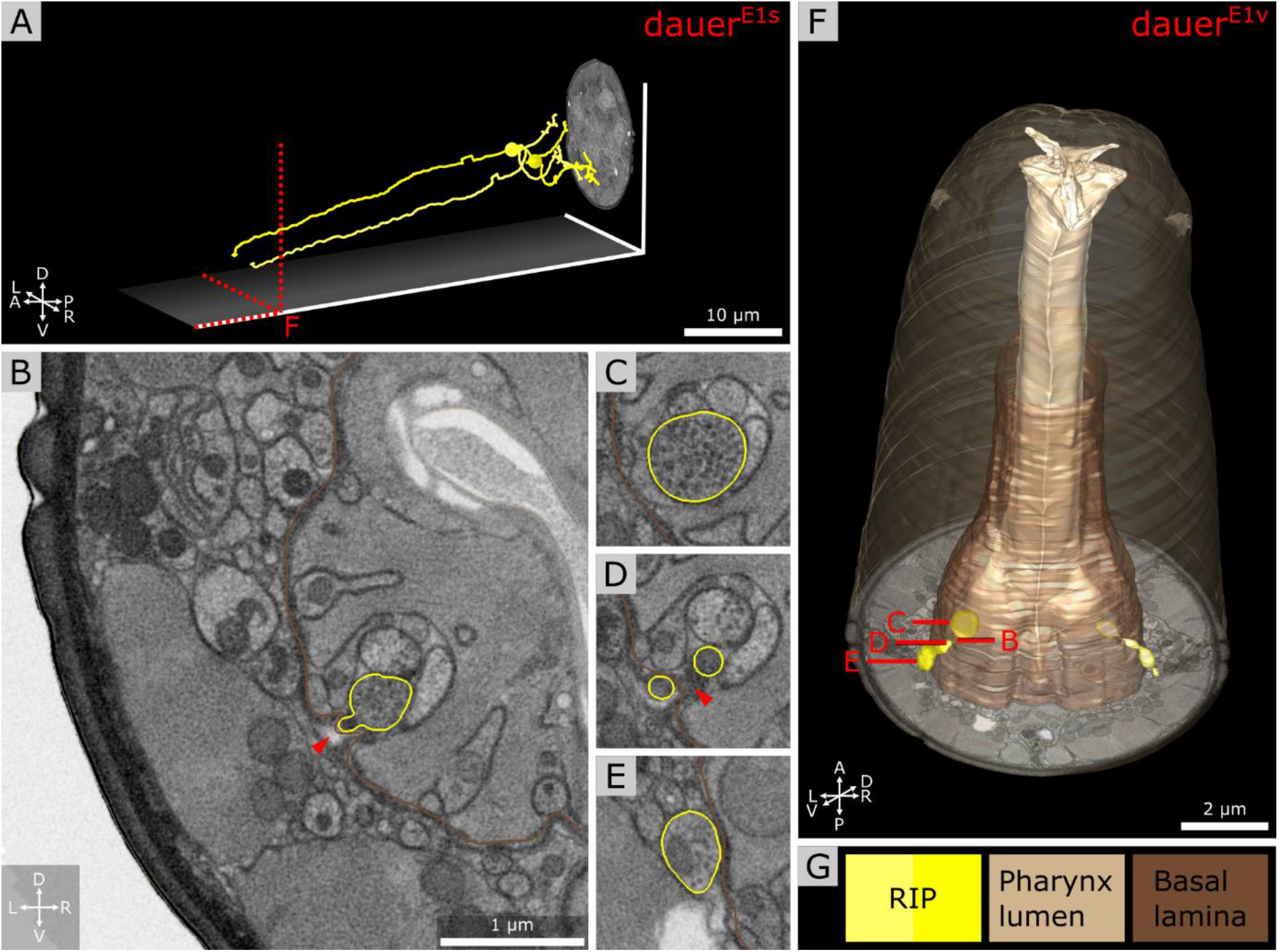
Anterior dendritic endings of RIP neurons of dauer enter the pharyngeal nervous system. **(A)** Tracing of RIP neurons. **(B)** Transverse FIB-SEM section at the position where the ending of RIPR is entering the pharyngeal nervous system through an opening of the pharyngeal basal lamina. **(C)** Bouton with electron dense vesicles of RIPR inside the pharyngeal nervous system. **(D)** RIPR split at the pharyngeal basal lamina just before its ending is entering the pharyngeal nervous system. **(E)** Transverse section posterior of **B** where RIPR forms a small bouton filled with electron dense vesicles. 3D reconstruction of RIP endings projecting through the pharyngeal basal lamina. **(G)** Names and color code of cells and structures shown.

## 4 Discussion

Earlier studies of IL sensilla cell morphologies of the dauer stage were limited by the ultrastructural preservation of classical fixation and the Z-resolution of ultramicrotomy sections (Albert and Riddle, 1983). To overcome these limitations, we used high-pressure freezing and FIB-SEM to reach a near-to-native preservation and high Z-resolution to identify precise dauer-specific features for the anterior sensory apparatus. Our findings confirm known and reveal new features that are consistent with an expansion of sensory function and plasticity as well as remarkable variability of sensory structures.

### 4.1 Morphological and functional implications of IL2 branching patterns

We confirm a distinct difference in the morphology between IL2L and IL2Q neuron branching types in dauer previously revealed by a light-microscopy study (Schroeder et al., 2013). These differences in their arborization pattern are mirrored at the molecular level by distinct expressions of transcription factors (Schroeder et al., 2013). Furthermore, these two neuron types differ in their synaptic connectivity in non-dauers (White et al., 1986; Witvliet et al., 2020). These differences suggest a functional difference between IL2L and IL2Q neurons that is morphologically manifested in dauer.

IL2 neurons are hypothesized to play roles in chemosensation as their endings are exposed to external environment in non-dauers (Ward et al., 1975; Ware et al., 1975; Lewis and Hodgkin, 1977; Albert and Riddle, 1983). In dauers, sensory endings of IL2 project less far anterior in relation to IL1 endings. This is opposite to non-dauers (Albert and Riddle, 1983). Potential retraction of IL2 sensory endings suggests that chemosensory functions in dauers may be partially replaced by increased mechanosensory or temperature perceptions mediated by dauer-extended branches important for nictation (Lee et al., 2011; Schroeder et al., 2013; Yang et al., 2020). Consistent with this idea, the *crown-like dendrites* at IL2L endings are located peripherally to other sensilla and are therefore in direct contact with the Hyp cell, in prime position to sense mechanical forces impacting on the anterior surface of the worm. Their location suggests sensing of nose touch. Furthermore, IL2QV’s *midline dendrites* (**Figure 4A**) shared the trajectory of AVM what contributes to gentle touch responses (Chalfie and Sulston, 1981), suggesting similar mechanosensitive functionalities for IL2Qs which are slightly different to IL2Ls. Previously, dauer IL2 neurons were proposed to play similar roles as mechano- and temperature-sensitive FLP neurons (Kaplan and Horvitz, 1993; Chatzigeorgiou and Schafer, 2011; Liu et al., 2012) in non-dauers, as they build dendritic branches in a similar receptive field (Albeg et al., 2011; Schroeder et al., 2013; Androwski et al., 2020). The morphological appearance of IL2L *crown-like dendrites* is fairly stable across dauers, with only slight differences in the 4° dendrites in one individual (**Supplementary Figure S2B-D**). It would be interesting to identify specific functional roles of the *crown-like dendrites*. Their overlap may indicate a signaling feedback.

### 4.2 Dauer-specific interactions of IL2Qs with other neurons

We found that dauer-specific dendritic endings of IL2Q reach the anterior projections of SAA or SAB (**Figure 4A**). SAA’s anterior projections originate from their axons (White et al., 1986) but are devoid of synapses (Witvliet et al., 2020). They are proposed to possess stretch receptors, and were modeled to be required for periodic bending of the anterior body (White et al., 1986; Sakata and Shingai, 2004; Vidal et al., 2015). An alternative model suggests that IL1 dendrites serve this role instead (Hart et al., 1995; Sakata and Shingai, 2004; Kindt et al., 2007). In non-dauers, anterior projections of SAB make neuromuscular junctions (Witvliet et al., 2020). It would be interesting to examine whether BWM cells receive synaptic input from SAB also in dauers, and whether SAB is required for nictation. Our current findings let us speculate that IL2Q may acquire mechanosensory functions and may regulate nictation via dauer-specific synapses with SAA or SAB (White et al., 1986; Witvliet et al., 2020).

### 4.3 Increased mechanosensation in dauers

Being a diapause state, dauer larvae exhibit remarkably quick responses and fast forward movement induced by mechanical stimuli (Cassada and Russell, 1975; Gaglia and Kenyon, 2009). Our hypotheses that dauer shows a higher degree of plasticity and is more mechanosensitive is further supported by finding a dauer-specific anterior extension of AVM dendrite in *Dauer^E1^*. AVM ends more posterior in non-dauers, approximately at the level of the pharyngeal metacorpus (Chalfie and Sulston, 1981; White et al., 1986). In the future, these observations should be further quantified by analyzing more individuals with imaging techniques that allow much larger sampling such as fluorescence microscopy with specific reporter lines (Ch’ng et al., 2003; Androwski et al., 2020).

### 4.4 ILLso-URX-AMso structures unique for dauers

Gas sensing neurons BAG interact with ILLso cells in *C. elegans* adults and ILLso are therefore hypothesized to provide structural stability (Doroquez et al., 2014; Cebul et al., 2020). Contrary to previous observations (Doroquez et al., 2014), gas sensing neurons URX interact with ILLso as well (Cebul et al., 2020). Similar observations were made for BAG and URX cell homologs in *P. pacificus* adults (Hong et al., 2019). We observed similar interaction in *C. elegans* dauers. ILQso do not form structures to physically interact with BAG or URX. Therefore, ILso cells also consist of two principally different categories similar to IL2 neurons.

It would be interesting to address if the morphological difference within ILso cells is also functionally relevant. ILso cells express receptors required for nose touch sensitivity mediated by IL1 (Hart et al., 1995; Kindt et al., 2007; Han et al., 2013). Such a sensitivity may be functionally further enhanced in dauer. Moreover, we discovered that in dauers the AMso cell encloses the URX neuron. In some cases, URX additionally enclose ILLso at the same position, forming a sandwich structure that was not observed in *C. elegans* (Doroquez et al., 2014) and *P. pacificus* (Hong et al., 2019) adults. Our investigation also revealed that AMso cells did not enclose URX neuron endings in the starved *L1^E4v^* larva (**Supplementary Figure S3E,F**). This is likely a dauer-specific adaptation that can be further investigated by examining the morphology of AMso and URX endings with fluorescent reporter lines previously described (Kim and Li, 2004; Low et al., 2019; Cebul et al., 2020; Fung et al., 2020).

We confirm that URX endings are ciliated in dauer (**Figure 5F,G**), as is the case in adults (Kazatskaya et al., 2020) and in *P. pacificus* (Hong et al., 2019). Surprisingly, we did not observe FLP endings to interact with ILLso or to be ciliated, as previously reported for non-dauers (Ward et al., 1975; Ware et al., 1975; Perkins et al., 1986; White et al., 1986; Doroquez et al., 2014) and dauers (Albert and Riddle, 1983).

### 4.5 Intracellular manifestations of dauer physiology

We observed convoluted whorls of membrane enclosed by or originating from either URX, BAG, AMso, ILLso, or Hyp cells (**Supplementary Figure S3A,B**). They resemble what has been described as exophers (Melentijevic et al., 2017). Exophers are formed by sensory neurons to overcome neurotoxic stress (Melentijevic et al., 2017). As dauers are in a constant state of physiological stress due to absence of food uptake (Cassada and Russell, 1975) it is not surprising to find exopher-like structures. Hyp cells are involved in cellular recycling of exophers (Melentijevic et al., 2017). As AMso and ILLso have hypodermal-like properties (White et al., 1986; Low et al., 2019; Cebul et al., 2020) and interact with URX or BAG, they could take on this task. Formation of exophers reinforces sensitivity of neurons (Melentijevic et al., 2017). These findings further support that sensory activity is increased in dauers.

### 4.6 Dauer specific adaptation and variability of amphid sensory arborizations

Carbon dioxide and temperature can be perceived by AFD neurons (Bretscher et al., 2011; Liu et al., 2012). Both parameters are important for dauers (Golden and Riddle, 1984; Hallem et al., 2011; Yang et al., 2020). We found a range between 84 and 112 microvilli per AFD neuron in dauers (**Table 1**), twice more than the previous estimation (Albert and Riddle, 1983). AWA terminal branches are between 12 and 24 in dauers (**Table 1**). Adults possess approximately 80 branches (Doroquez et al., 2014), implicating a developmental increase as reported for FLP (Albeg et al., 2011). We observed a wing-like swelling in one AWA branch in *L1^E4v^*. This may be an effect of starvation as similar starvation-dependent morphological changes for AWB neurons in adults have been described (Mukhopadhyay et al., 2008). In non-dauers, AWC are rather T-shaped (Ward et al., 1975; Doroquez et al., 2014). In dauers, we found more a Y-shape, and the radial span of AWC wings in our data sets are larger (up to 343°, **Table 1**) than previously reported (Albert and Riddle, 1983).

Morphological differences in sensory structures between individuals may be triggered by the external environment or intrinsic influences. In contrast to *developmental plasticity*, where the relevance of environmental influences is clear, we here prefer to use the term *variability*, as the environmental impact has not been investigated in detail yet. There is variability in morphology of amphid sensory endings among dauers. AWC in *Dauer^E2^* showed cupola-shaped structures in parts of the wing. Branch direction of AWBL in *Dauer^E3^* was different from others. We hypothesize that phenotypic variability among dauers might be a preadaptation for the dauer population to find and adapt to new habitats.

### 4.7 Interactions between the pharyngeal and somatic nervous system

Pharyngeal pumping is inactivated in dauers (Cassada and Russell, 1975; Albert and Riddle, 1983). We found that RIP, the only direct neuronal connection between pharyngeal and somatic nervous system, project in a similar manner as in non-dauers (Ward et al., 1975). RIP projection terminals contain synaptic varicosities filled with electron dense vesicles in our observations which often contain neuropeptides (Ann et al., 1997; Salio et al., 2006). Similar varicosities were not found in starved *L1^E4^* and have not been described in other non-dauers (Ward et al., 1975; Albertson and Thomson, 1976). In dauers, RIP endings may also function as motor neurons (**Supplementary Figure S5B**).

In non-dauers, RIP regulates inhibition of pharynx pumping triggered by light touch perceived by other neurons (Chalfie et al., 1985; Avery and Thomas, 1997). RIP could be responsible for pharynx inhibition in dauers. As IL1 are mechanosensitive (Hart et al., 1995; Kindt et al., 2007) and IL2 are hypothesized to be so with high activity, both may dauer-specifically enhance inhibition of the pharynx via RIP similarly (White et al., 1986).

In summary, we observed expansion, plasticity, and variability of the sensory neuron apparatus in dauers. An enhancement of sensory perception facilitates finding of more favorable environments to re-enter the reproductive life cycle.

## 5 Author Contributions

All authors had full access to all the data in the study and take responsibility for the integrity of the data and the accuracy of the data analysis. YS, PK, MZ, CS: study concept and design; SB, DW, SMM, ST, BM, AMS: data analysis and interpretation; SB, CS, MZ: drafting of the manuscript; all authors: critical revision of the manuscript for important intellectual content; SMM, AMS: acquisition of data; MZ, CS: study supervision.

## 7 Acknowledgments

We thank for the support from co-annotators F. Schinke, A. Hutchings, A. Gerber, J. Olbrich, C. Sommer, P. Dinkel, J. Karl, and R. Jacobalis. We thank D. Mastronarde, J. Bush, and S. Spielhaupter, for support and helpful hints about 3dmod software. We thank M. Wang, D. Fon, and A. D.T. Samuel for CATMAID server support. We thank D. Bunsen, C. Gehrig-Höhn, B. Trost, and N. Schieber for excellent technical support and helpful hints regarding sample preparation. We thank the electron microscopy core unit at the Max Planck Institute of Experimental Medicine for access to their Crossbeam 540.

## 8 Supplementary Material

See supplementary material document and videos.

