## Supplementary Document for "Structural analysis of the *C. elegans* dauer larval anterior sensilla by Focused Ion Beam-Scanning Electron Microscopy"

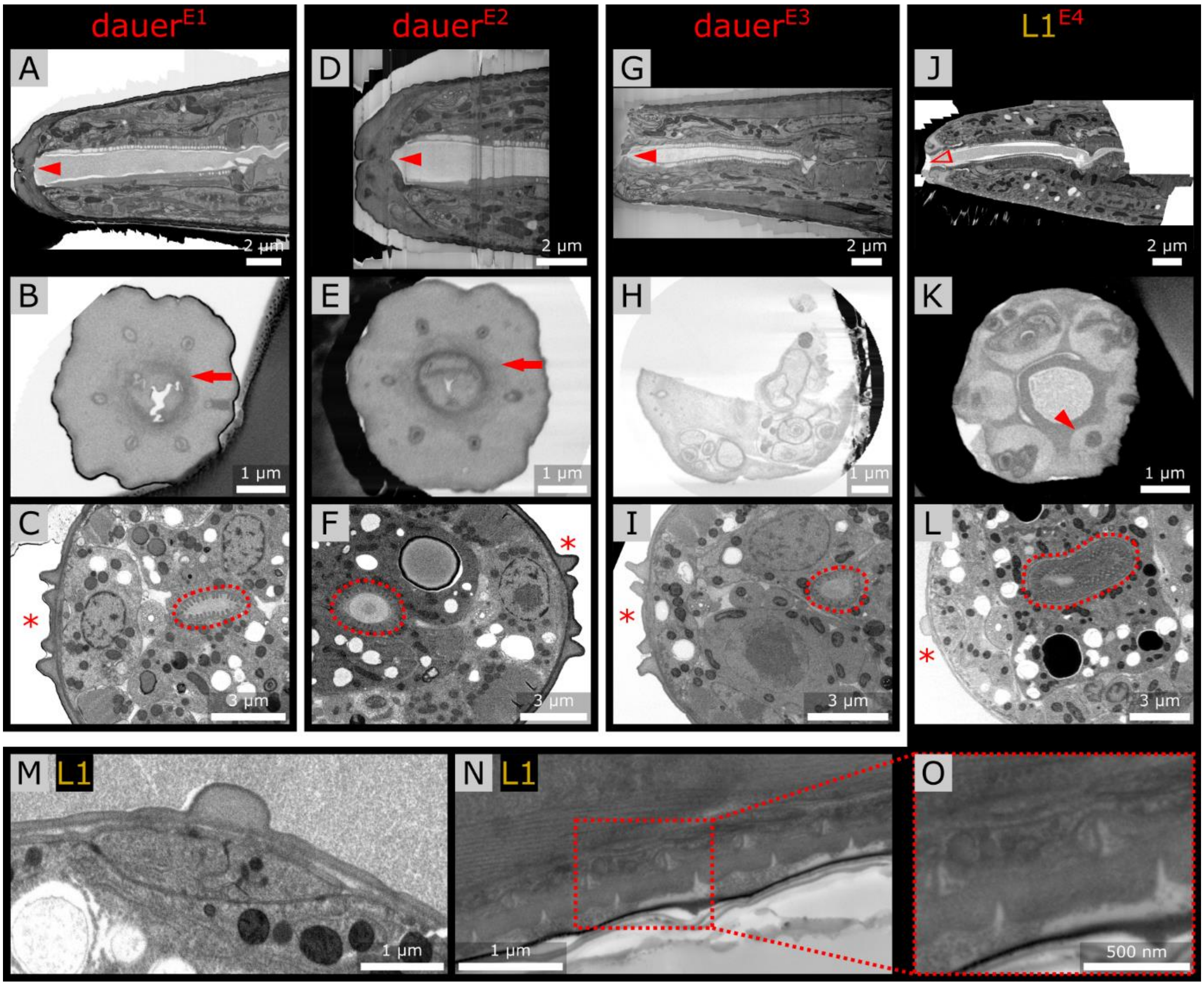


**Figure S1.** Anatomical features of the investigated dauer larvae and L1 larva. **(A)** Longitudinal virtual section through the volumetric reconstructed image stack of the first dauer larva. The mouth is closed (arrowhead). **(B)** Transverse section of the anterior worm tip showing dauer-specific cuticle fibers (arrow). The nose appears flat as the six lips are not clearly constricted compared to the L1 larva in **K**. **(C)** Transverse section in the mid-region of the worm showing reduced intestinal structures (dashed outline) and alae (asterisk) in shape specific for dauer larvae. **(D-F)** Second dauer shown in the same manner as the first dauer larva in **A-C**. **(G-I)** Third dauer shown in the same manner as the first dauer larva in **A-C**. **(G,H)** The anterior tip is partly missing. **(J-L)** L1 larva shown in the same manner as the first dauer larva in **A-C**. Note that the mouth is open (**J**, non-filled arrowhead). The six lips are more prominently present, here visible for one lip curved on the side facing the mouth opening (**K**, arrowhead), the intestinal structures are more expanded (**L**, dashed outline), and the alae of the L1 larva (**L**, asterisk) differ in shape from those of dauer. **(M)** Alae of the L1 larva higher magnified. Note that there are two cuticles. The alae have one big ridge arising from the upper cuticle. **(N)** SEM image of longitudinal ultramicrotome section of the L1 larva focusing on the cuticle. The worm started to produce the next cuticle. **(O)** Detail of **N**.

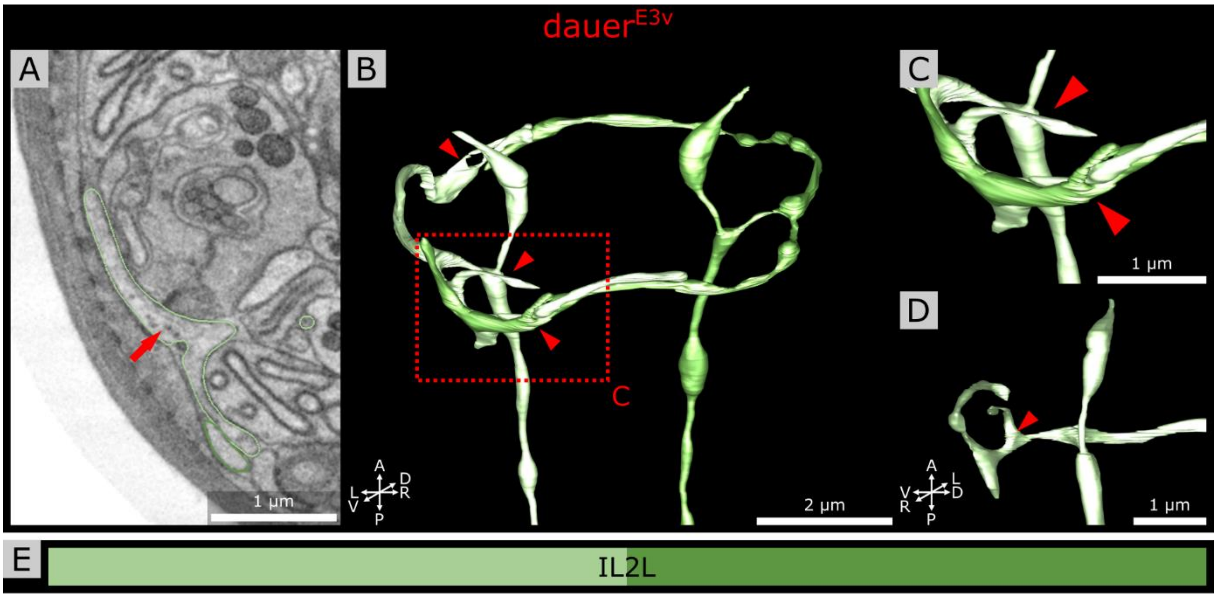


**Figure S2.** *Crown-like dendrites* of IL2L neurons in another dauer data set showing individual differences. **(A)** Transverse FIB-SEM sections through the 3° dendrite of IL2LL of *Dauer$^{E3v}$*. A few electron dense vesicular structures are visible in the *crown-like dendrite* (arrow). **(B)** Volumetric reconstruction of the anterior IL2L neuron endings of *Dauer$^{E3v}$.* Crown-like dendrites of both IL2L neurons extend 4° dendrites (arrowheads). **(C)** 4° dendrites shown in **B** higher magnified. **(D)** One 4° dendrite (arrowhead) of IL2LL from another perspective. **(E)** Names and color code of cells shown.

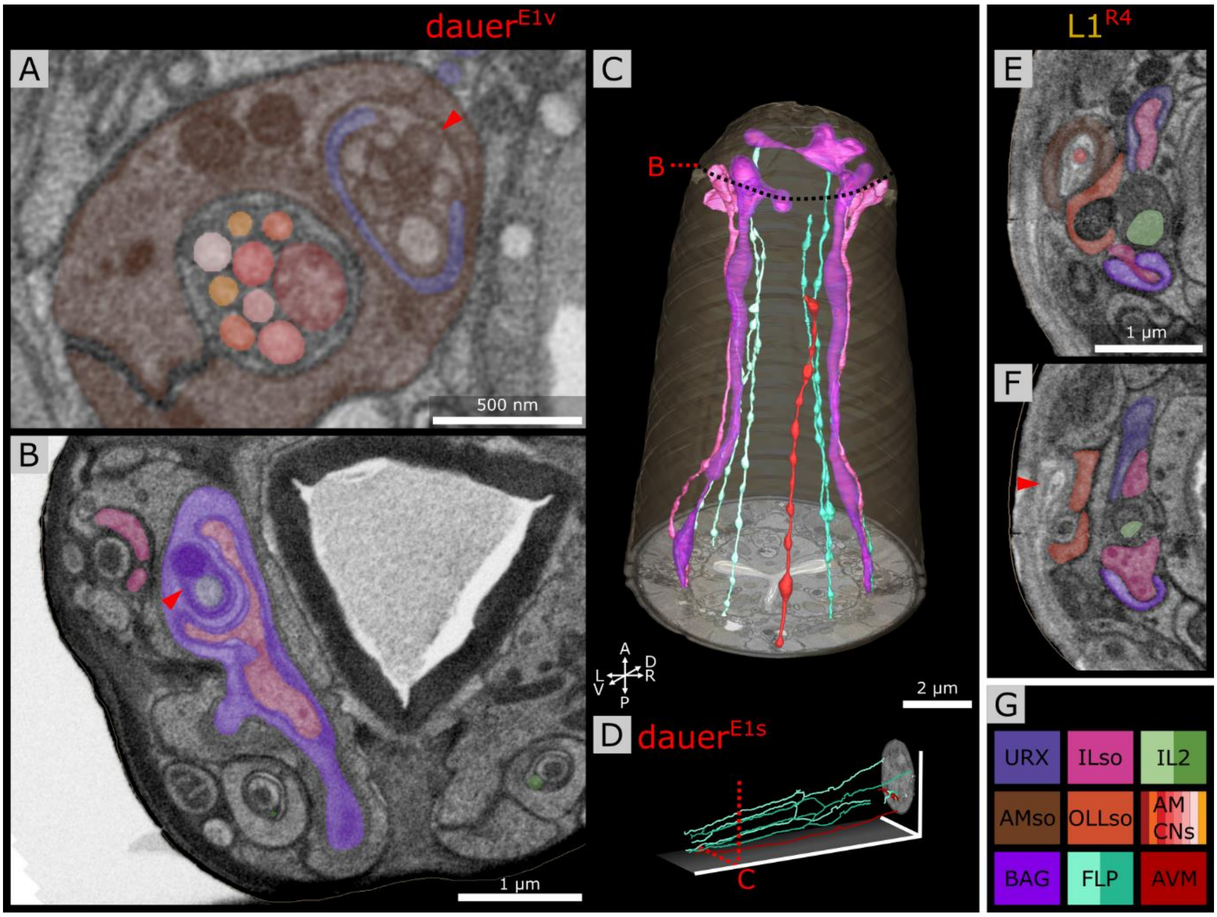


**Figure S3.** Further investigation of ILLso and AMso cell endings as well as non-sensilla neurons. **(A)** URXL is enclosed by AMsoL cell and encloses big vesicular structures (arrowhead). **(B)** BAGL ending encloses one branch of ILLsoL. BAGL also encloses two vesicular structures (arrowhead) of which one is electron dense and the other might originate from URXL. **(C)** Volumetric 3D reconstruction of ILsoL cells with two branches each of which one is enclosed by BAG endings while FLP do not have any branch or ILso-interaction. AVM reaches almost into the anterior sensilla region. **(D)** Tracing of FLP neurons and the AVM neuron. **(E)** Transverse section of ILLsoL ending in an L1 larva. ILLsoL has two branches which are enclosed by URXL and BAGL. URXL is not enclosed by AMsoL cell. **(F)** OLLsoL cell encloses the amphid opening which is in this case formed by respective Hyp cell (arrowhead). **(G)** Names and color code of shown cells.

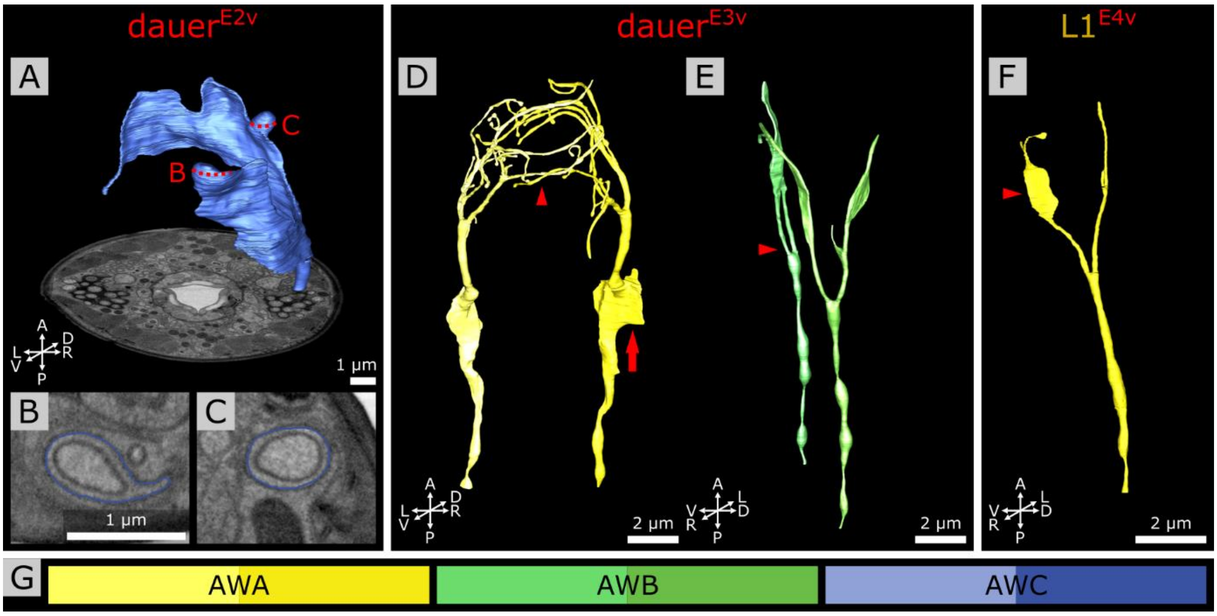


**Figure S4.** Individual findings of amphid wing neuron endings in dauer larvae and one L1 larva. **(A)** Volumetric reconstruction of the AWCR ending of *Dauer*$^{E2}$ fusing twice with itself (dashed lines) to a capsule shape. **(B,C)** Transverse sections through the AWCL capsules from **A**. **(D)** Volumetric reconstruction of AWA neuron endings of *Dauer*$^{E3}$ which are widely intertwining (arrowhead). AWAR shows a wing-like thickening at the cilia base (arrow). **(E)** Volumetric reconstruction of AWB neuron endings. The two AWBL branches both project into the ventral side (arrowhead). **(F)** Volumetric reconstruction of AWAR neuron ending of *L1*$^{R4}$ which has a wing-like morphology at one branch (arrowhead). **(G)** Names and color code of shown cells.

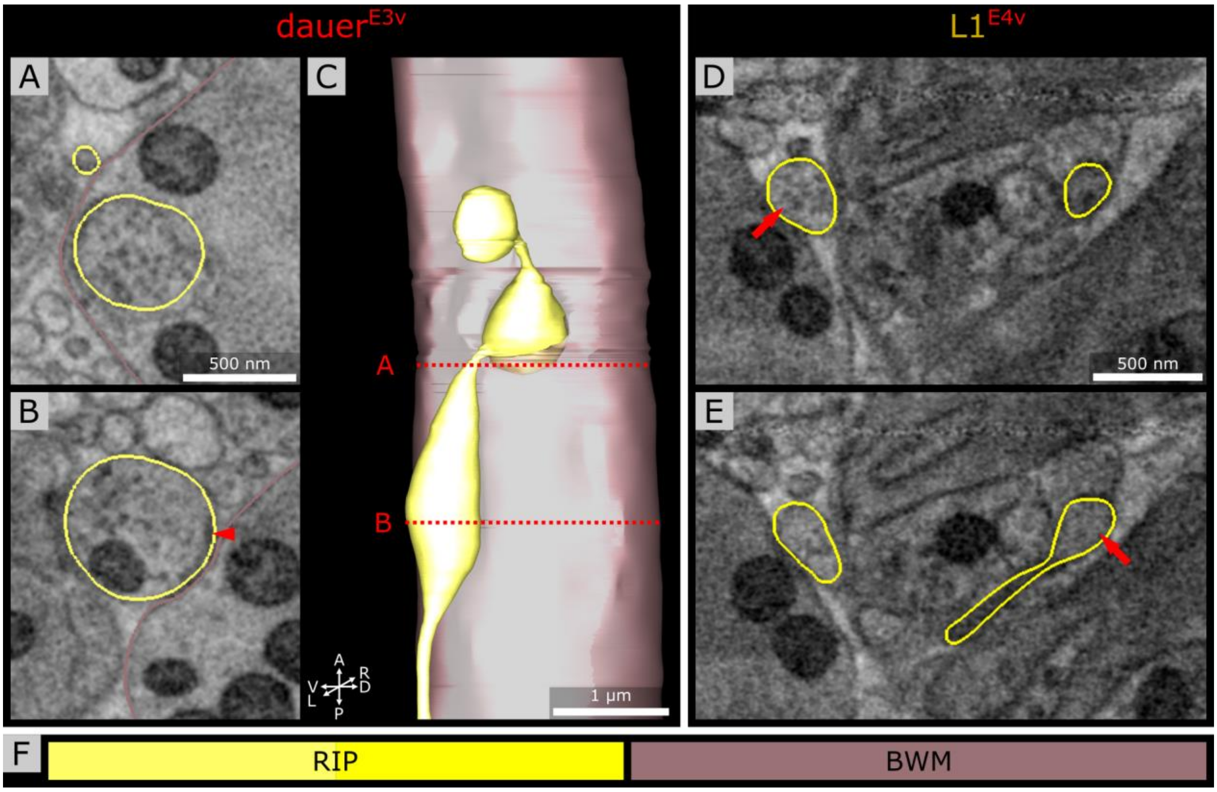


**Figure S5.** RIP anterior endings enter the pharyngeal nervous system in another dauer individual and in one L1 larva. **(A)** Section of the RIPL ending enclosed by BWM cell and filled with electron dense vesicles. **(B)** Section of RIPL ending showing an electron dense projection facing the respective BWM cell (arrowhead). **(C)** Volumetric reconstruction of the RIPL ending. RIPL has a big bouton enclosed by respective BWM cell (section in **A**) before its ending is entering the pharynx. The ending inside the pharyngeal nervous system has a bouton-like structure as well. **(D)** Section of RIPR ending of an L1 larva showing a bouton with electron dense vesicles (arrow) before entering the pharyngeal nervous system like in dauer but to a lesser extent. **(E)** Section of RIPR ending inside the pharyngeal nervous system just passing the pharyngeal basal lamina with a bouton-like structure (arrow) but again to a lesser extent compared to dauer.

**Video S1.** Volumetric reconstruction of the cell endings at the anterior tip of a *C. elegans* dauer larva.

See Supplementary Material.

**Video S2.** Dendritic branches of the IL2Q neurons in one dauer individual in the context of the complete image stack and volumetric reconstruction of the anterior worm cell endings.

See Supplementary Material.

**Table S1.** Stage specific anatomical features for identification of dauer and L1 stage under starving conditions.

| Determination | Existence or shape of feature | Data set | |
|---|---|---|---|
| | | E1-3 | E4 |
| **Stage** | **Cuticle** | Not investigated. | Next cuticle layer formation in process, indicating active development (Popham and Webster, 1979) and the analyzed larva stage to be just before its end. Excluding developmental arrest. See **Supplementary Figure S1M-O**. |
| | **Alae** | Existing, like known dauer-specific shape (Cassada and Russell, 1975; Singh and Sulston, 1978; Cox et al., 1981; Albert and Riddle, 1983). See **Supplementary Figure S1C,F,I**. | Existing, alae on the outer cuticle layer similar to known L1-specific shape (Cox et al., 1981) and like shape with second cuticle (Popham and Webster, 1979). Excluding L2d-dauer molt as dauer-specific alae would be on the inner cuticle (Singh and Sulston, 1978). See **Supplementary Figure S1L,M**. |
| | **Mouth opening** | Closed, like known for dauer and L2d-dauer molt (Albert and Riddle, 1983; Golden and Riddle, 1984). See **Supplementary Figure S1A,D,G**. | Open, known for L1 (Baugh and Sternberg, 2006; Baugh et al., 2009) and excluding L2d-dauer molt as it would be closed (Golden and Riddle, 1984). See **Supplementary Figure S1J**. |
| | **Lips** | Less distinct, like known for dauer which let the *C. elegans* nose appear flatter than that of L2 stages (Albert and Riddle, 1983). Not clearly observable in data set E3 as structure is partly broken. See **Supplementary Figure S1B,E,H**. | More prominent lips than the other larvae. See **Supplementary Figure S1K**. |
| | **Cuticular fibers** | Existing, like known for dauer around their mouth opening (Albert and Riddle, 1983). Not observable in data set E3 as structure is partly broken. See **Supplementary Figure S1B,E,H**. | Not existing, excluding dauer as it was only observed in dauers up to now (Albert and Riddle, 1983). See **Supplementary Figure S1K**. |
| | **Intestine** | Reduced, like known for dauer (Popham and Webster, 1979). See **Supplementary Figure S1C,F,I**. | Not reduced, similar to known shape of L1 larva (Popham and Webster, 1979; Maduro, 2017). See **Supplementary Figure S1L**. |

…

…

| Determination | Existence or shape of feature | Data set | |
|---|---|---|---|
| | | **E1-3** | **E4** |
| **Stage** | **Length** | 435 µm measured for E3, similar known for dauer (Cassada and Russell, 1975; Cox et al., 1981). Measurement only possible for E3. | Measurement not possible. |
| | **Diameter** | ~14-18 µm measured in the body middle segment, similar known for dauer (Cassada and Russell, 1975; Cox et al., 1981). | ~17 µm measured in the body middle segment, similar known for L1 (Cassada and Russell, 1975; Cox et al., 1981). |
| **Biological sex** | **CEM neurons** | Not existing, excluding male as it is male-specific (Sulston et al., 1983). | Not existing, excluding male as it is male-specific (Sulston et al., 1983). |
| **Result** | | **Stage: Dauers**<br>**Sex: Hermaphrodites** | **Stage: L1**<br>**Sex: Hermaphrodite** |

**Table S2.** Properties of acquired FIB-SEM data sets.

| Data set | Image data | Body segment | Direction of acquisition |
|---|---|---|---|
| ***Dauer***$^{E1}$ | 82.312 µm with 10,289 image slices. Voxel size: 5 x 5 x 8 nm. Note: data set uploaded to CATMAID contains six images more. | From the amphid commissure to the anterior worm end. | Transversal from posterior to anterior. |
| ***Dauer***$^{E2}$ | 7.768 µm with 971 image slices. Voxel size: 5 x 5 x 8 nm. | From the Z-level where microvilli of AFD neurons are visible to the anterior worm end. | Transversal from posterior to anterior. |
| ***Dauer***$^{E3}$ | 18.4 µm with 2,300 image slices. Voxel size: 5 x 5 x 8 nm. | From the Z-level posterior to where the amphid neurons just are getting enclosed by the AMsh cell to the anterior worm end. | Transversal from posterior to anterior. |
| ***L1***$^{E4}$ | 18.770 µm with 3,754 image slices after stack transformation. Voxel size: 5 x 5 x 5 nm. | Anterior worm end. | Longitudinal from ventral to dorsal. |

### Figure and video creation

All figures were arranged with the software Inkscape (Inkscape.org Team, 2021). Schemes were created as vector graphics with Inkscape on basis of pixel snapshots from contour drawings from 3dmod (Kremer et al., 1996) reconstruction or vector graphic snapshots from skeleton tracing from CATMAID (Saalfeld et al., 2009). 2D labels from 3dmod were merged with respective EM images in GIMP as separated 3dmod pixel snapshots (GIMP.org Team, 2021). Brightness and contrast of EM images in 3D view of 3dmod were post-adjusted with 3dmod. Snapshots of **Supplementary Figure S1D,G** were post-denoised with VSNR plugin in Fiji and once more brightness and contrast re-adjusted in 3dmod. 3dmod was also used to adjust images shown in 3D view of CATMAID. After adjustment, images were then merged with the tracing skeleton vector graphics in Inkscape. These EM images were also rendered manually by blacking pixels on the outside around the worm with the software GIMP. For **Figure 1B**, EM images were rendered by creating as mask from the outline of the body wall cuticle from 3dmod and subtracting it from the image stack in Fiji (Schindelin et al., 2012). This rendered image stack was also taken for **Supplementary Videos S1**, **S2**. Image sequences for videos were created with 3dmod. Video labels were created with Inkscape. Labels and snapshots were merged with the Python Imaging Library Pillow (Python-Pillow.org Team, 2021) and exported as video with the OpenCV-Python library (opencv-python Team, 2021). For **Supplementary Video S2**, tracing skeletons from CATMAID were transformed into a 3dmod model. For that, coordinates of tracing nodes were exported as .csv file from the 3D view of CATMAID, prepared, and then imported into 3dmod with the *mod2point* function.

### References Supplement


Albert, P. S., and Riddle, D. L. (1983). Developmental alterations in sensory neuroanatomy of the Caenorhabditis elegans dauer larva. *J. Comp. Neurol.* 219, 461–481. doi:10.1002/cne.902190407.

Baugh, L. R., DeModena, J., and Sternberg, P. W. (2009). RNA Pol II Accumulates at Promoters of Growth Genes During Developmental Arrest. *Science* 324, 92–94. doi:10.1126/science.1169628.

Baugh, L. R., and Sternberg, P. W. (2006). DAF-16/FOXO Regulates Transcription of cki-1/Cip/Kip and Repression of lin-4 during C. elegans L1 Arrest. *Curr. Biol.* 16, 780–785. doi:10.1016/j.cub.2006.03.021.

Cassada, R. C., and Russell, R. L. (1975). The dauerlarva, a post-embryonic developmental variant of the nematode Caenorhabditis elegans. *Dev. Biol.* 46, 326–342.

Cox, G. N., Staprans, S., and Edgar, R. S. (1981). The cuticle of Caenorhabditis elegans: II. Stage-specific changes in ultrastructure and protein composition during postembryonic development. *Dev. Biol.* 86, 456–470. doi:10.1016/0012-1606(81)90204-9.

GIMP.org Team (2021). *GIMP Version 2.10.24*. Available at: https://www.gimp.org/ [Accessed June 13, 2021].

Golden, J. W., and Riddle, D. L. (1984). The Caenorhabditis elegans dauer larva: Developmental effects of pheromone, food, and temperature. *Dev. Biol.* 102, 368–378. doi:10.1016/0012-1606(84)90201-X.

Inkscape.org Team (2021). *Inkscape Version 1.1*. Available at: https://inkscape.org/ [Accessed June 13, 2021].

Kremer, J. R., Mastronarde, D. N., and McIntosh, J. R. (1996). Computer visualization of three-dimensional image data using IMOD. *J. Struct. Biol.* 116, 71–76. doi:10.1006/jsbi.1996.0013.

Maduro, M. F. (2017). Gut development in C. elegans. *Semin. Cell Dev. Biol.* 66, 3–11. doi:10.1016/j.semcdb.2017.01.001.

opencv-python Team (2021). *opencv-python*. OpenCV Available at: https://github.com/opencv/opencv-python [Accessed June 13, 2021].

Popham, J. D., and Webster, J. M. (1979). Aspects of the fine structure of the dauer larva of the nematode Caenorhabditis elegans. *Can. J. Zool.* 57, 794–800. doi:10.1139/z79-098.

Python-Pillow.org Team (2021). *Python Pillow*. Available at: https://python-pillow.org/ [Accessed June 13, 2021].

Saalfeld, S., Cardona, A., Hartenstein, V., and Tomančák, P. (2009). CATMAID: collaborative annotation toolkit for massive amounts of image data. *Bioinformatics* 25, 1984–1986. doi:10.1093/bioinformatics/btp266.

Schindelin, J., Arganda-Carreras, I., Frise, E., Kaynig, V., Longair, M., Pietzsch, T., et al. (2012). Fiji: an open-source platform for biological-image analysis. *Nat. Methods* 9, 676–682. doi:10.1038/nmeth.2019.

Singh, R. N., and Sulston, J. E. (1978). Some Observations On Moulting in Caenorhabditis Elegans. *Nematologica* 24, 63–71. doi:10.1163/187529278X00074.

Sulston, J. E., Schierenberg, E., White, J. G., and Thomson, J. N. (1983). The embryonic cell lineage of the nematode Caenorhabditis elegans. *Dev. Biol.* 100, 64–119. doi:10.1016/0012-1606(83)90201-4.
